# Muscle contraction as a Markov Process - II: x-ray interference data (M3 & M6 reflections) imply myosin cross-bridge motions are controlled by structural transitions along actin filaments

**DOI:** 10.1101/2021.01.16.426939

**Authors:** Clarence E Schutt, Vladimir Gelfand, Eli Paster

**Affiliations:** Department of Chemistry, Princeton University

**Keywords:** muscle contraction theory, independent force generators, actin filaments, Huxley-Simmons, x-ray fiber patterns, quick release, myosin cross-bridges

## Abstract

The unit underlying the construction and functioning of muscle fibers is the sarcomere. Tension develops in fibers as thousands of sarcomeres arranged in series contract in unison. Shortening is due to the sliding of actin thin filaments along antiparallel arrays of myosin thick filaments. Remarkably, myosin catalytic heads situated across the center M-line of a sarcomere are separated by a distance that is a half integral of the 14.5 nm spacing between successive layers of myosin heads on the thick filaments. This results in the splitting of the 14.5 nm meridional reflection in X-ray diffraction patterns of muscle fibers. Following a quick drop in tension, changes in the relative intensities of the split meridional peaks provide a sensitive measure of myosin head movements. We use published data obtained with the x-ray interference method to validate a theory of muscle contraction in which cooperative structural transitions along force-generating actin filaments regulate the binding of myosin heads. The probability that an actin-bound myosin head will detach is represented here by a statistical function that yields a length-tension curve consistent with classical descriptions of the recovery of contracting muscle fibers subjected to millisecond drops in tension.

## Introduction

It is widely supposed that contraction of skeletal muscle is powered by myosin S1 motor domains projecting from myosin thick filament shafts pulling on passive actin filaments as sarcomeres shorten (Huxley, 1969; Huxley & Simmons, 1971; Holmes, 1997; Cooke, 1986). The free energy for this process is provided by the hydrolysis of ATP on myosin S1 domains attaching and detaching as actin filaments move along them (Lymn & Taylor, 1971). Five decades of biochemical, structural and biophysical research have lent considerable support to this hypothesis (Geeves & Holmes, 2005), yet problems remain in reconciling *in vitro* studies on isolated components of the system (Harada et al., 1990; Higuchi & Goldman, 1991; Higuchi & Goldman, 1995); Toyoshima et al., 1990; Uyeda et al., 1990, particularly with regard to the timing of phosphate release by muscle fibers generating forces (Månsson et al., 2015, Schutt & Lindberg, 1998), and in assigning distinct structural states to cross-bridge dynamics (Pollack,1990; Morel, 2016).

A long-standing goal of muscle research is to explain changes in sarcomere length in terms of the motions of myosin, actin and tropomyosin during a contraction. The mechanism of force generation can be studied by rapid release of actively contracting striated muscle fibers and observing the transient recovery of tension (Podolsky, 1960; Podolsky & Nolan, 1969; Huxley & Simmons, 1971; Ford *et al*., 1977; Dobbie et al., 1998). In a pioneering series of x-ray experiments, H.E. Huxley and colleagues (Huxley & Brown, 1967; Huxley et al, 1983; Xu et al., 1987; Huxley et al., 1994) demonstrated that millisecond time resolution is obtainable from muscle fibers during recovery from a quick drop in tension. Subsequent technical advances in X-ray flux production and electronic data collection have enabled the extension of these experiments to include simultaneous measurement of large regions of x-ray patterns from single fibers with a time resolution approaching 10 μs. One of the most striking discoveries made with these experimental arrangements is the splitting of the myosin-based (~14.5 nm) meridional reflection (the M3) into two peaks owing to interference between the two arrays of myosin heads across the center of a sarcomere (Juanhuix *et al*., 2001; Linari *et al*., 2000). These data can be used to measure movements of myosin heads attached to actin with sub-nanometer precision in contracting muscle fibers (Piazzesi et al., 2002; 2004) and whole muscle cells (Huxley et al., 2006) during quick releases.

These observations can be used to test the predictions of an alternative theory of muscle contraction based on the propagation of localized length changes in actin filaments as they move towards the center of the sarcomere (Schutt & Lindberg, 1992, Schutt & Lindberg, 1993; Schutt et al., 1995a; Schutt & Lindberg, 1998). The concept of an actin power-stroke emerged from x-ray studies on profilin:β-actin crystals where actin monomers along the 7.15 nm two-fold screw b-axis of the unit cell are organized into ribbons with strong inter-subunit bonds reminiscent of those seen in biologically-relevant fibers such as sickle-cell hemoglobin (Schutt et al., 1989; Schutt et al.,1993; Chik et al., 1996). The impetus for a new theory was the realization that 12 actin subunits in a ‘ribbon segment’ are commensurate with the repeat distance (42.9 nm) of myosin heads projecting from thick filaments (Huxley and Brown, 1967). Notably, four neighboring actin subunits in a ribbon segment centered on the trigonal position in myosin thick filament lattice can span a distance of 14.3 nm, enabling S1-domains from different thick filaments to be bound simultaneously to opposite sides of a twisted ribbon. This metastable intermediate would have a quasi two-fold screw axis that would contribute scattering to the second order peak of the 7.16 nm reflection, usually referred to as the M6.

In the actin power-stroke model, crawling actin filaments are activated by their interactions with myosin cross-bridges, which initiate local length changes in actin subunits, *accompanied by the uptake of ATP on actin.* The thin filaments generate local forces (100 pN) as they contract, utilizing the Gibbs free energy from ATP hydrolysis *on actin* to produce work, by taking purchase on bound myosin crossbridges and pulling on tropomyosin (Schutt and Lindberg, 1998). At any given moment only a small number of actin subunits are contracting along a thin filament, but as each one completes its contraction it triggers its neighbor to shorten thereby maintaining a constant force pulling on the associated tropomyosin filament. The axial and azimuthal motions of *attached* myosin cross-bridges in this model are passive and arise from forces acting on them by actin filaments crawling towards the center of the sarcomere. During a quick release of tension, the elastic cross-bridges return to unstrained actin-bound positions, only to be re-bent and twisted during tension redevelopment by actin filaments undergoing ribbon-to-helix transitions.

We have previously shown that this theory meets the requirements of A.F. Huxley’s (Huxley, 1980) independent force generator hypothesis (Schutt and Lindberg, 1992), given that tropomyosin filaments sum the forces developed by local contractions of actin subunits. This theory also explains how cooperative transitions in actin can control myosin on-off rates on thin filaments while yielding self-consistent free energy relations that account for Hill’s force-velocity curve (Hill, 1938) and the Fenn Effect (Schutt and Lindberg, 1998). Here we provide a detailed model for calculating the numbers and positions of bound myosin heads during tension recovery following a quick release in experiments of the Huxley-Simmons type and illustrate how to implement this model to account for data obtained with x-ray interference techniques (Piazzesi et al, 2002; Huxley et al. 2006; Reconditi et al., 2011). The axial movements (in the direction of shortening) of myosin cross-bridges inferred from these measurements are unexpectedly small given the assumptions of the rotating myosin cross-bridge theory of contraction (Piazzesi *et al*., 2002). In the theory presented here, actin-induced azimuthal rotations (about the thick filament axis) take up part of the strain and provide an explanation for the unexpected behavior of the M3 and M6 x-ray reflections (1^st^ and 2^nd^ orders of the myosin 14.3 nm reflection).

### The Huxley-Simons Formalism

Huxley & Simmons (1971) reported that during a sudden length change tension in an isometrically contracting muscle fiber tension drops instantaneously, followed by a complex recovery consisting of several phases, where elastic elements are stretched again in the T_1_ phase, followed by partial recovery of tension (T_2_ phase) within a few milliseconds, with the isometric value reached in the later (T_3_ and T_4_) phases on slower time scales. The recovery of tension during the T_2_ phase is exponential:

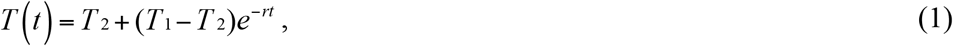

where *r* is the rate of tension recovery (Huxley & Simmons, 1971). The rate r depends drastically on the value of the applied length step, and is approximated well by:

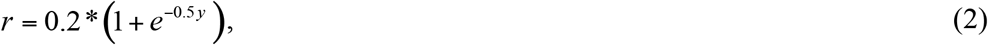

where y is the magnitude of the length step (positive for stretch).

Two structural elements must be present in series in order to explain these observations (Huxley & Simmons, 1971). One of them is an elastic element that responds instantaneously to the change in tension, giving rise to the T_1_ curve. The other, the active force generator, is an element with viscous as well as elastic properties, which begins to regenerate tension (T_2_ curve) following a delay of several milliseconds. Huxley and Simmons suggested that the elastic and force-generating elements both reside in myosin cross-bridges; specifically, the force-generating element was identified with the catalytic myosin heads (S1), and the elastic element was placed in the S2 links. In order to explain the T_2_ curve, they suggested that cross-bridges occupy any of a number of bound stable states, each having a different potential energy. The total potential energy of an attached cross-bridge is the energy of the bound state plus the potential energy involved in stretching the elastic S2 link. Following a length change, the crossbridges gradually redistribute amongst the bound states according to Boltzmann statistics in order to minimize the total potential energy. This model attributes quick tension recovery to the tendency of myosin heads to rotate to positions of lower potential energy. The rate of tension recovery reflects the finite speed of movement of the system from one stable set of positions position to another.

Huxley and Simmons (1971) presented a simplified quantitative model, based on only two stable states, and derived an expression for the recovered tension as a function of length step:

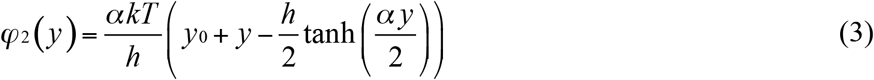

Here ϕ_2_ is tension in the T_2_ phase, y_0_ is the average pre-release extension of S2 links, y is the length step (positive for stretch), h is the increase of the S2 length when a cross-bridge shifts from state 1 to state 2, *a* is a parameter (which was taken as 5*10^8^ m^-1^), k is the Boltzmann constant, T is temperature. We will demonstrate below that this equation, which accounts for all of the features of the T_2_ length-tension curve, can be derived from an independent force generator model based on cooperative structural transitions in actin..

### Problems posed by x-ray interference measurements on muscle

One of the assumptions of the original Huxley-Simmons (1971) model is virtual inextensibility of the myosin thick and actin thin filaments. The discovery of significant filament elasticity (Huxley *et al*., 1994; Kojima *et al*., 1994; Wakabayashi *et al*., 1994; Higuchi *et al*., 1995) has altered the view on where the classical elastic element is located. It is widely accepted that the instantaneous elasticity is distributed between the myosin heads and the thick and thin filaments. Attributing the force-generating element to myosin heads still remains a characteristic feature of all models based on the cross-bridge theory. Muscle fibers are able to regenerate tension when consecutive steps, even as short as 8 ms apart are applied during the T_2_ phase of tension recovery of a previous step (Lombardi *et al*., 1992; Irving *et al*., 1992). In order to explain this phenomenon, the rapid detachment-attachment hypothesis (Brenner, 1991; Lombardi *et al*., 1992, 1995) was proposed, which asserts that myosin heads exert multiple power-strokes for each ATP molecule consumed, implying that the free energy of ATP hydrolysis can be parceled out during several relatively short power-strokes applied at different points along the sliding actin filament, an explanation that has failed to garner support (Schutt & Lindberg, 1998; Piazzesi et al, 2002).

The M3 reflection corresponding to the axial spacing (~14.5 nm) of myosin heads and its second order reflection the M6 (~7.25 nm) are split into two distinct peaks (I_HA_ and I_LA_, higher and lower angle respectively), with a prominent shoulder at ~14.3 nm. The appearance of split reflections can be explained on the basis of X-ray interference (Juanhuix *et al*., 2001; Linari *et al*., 2000) between waves scattered by arrays of 49 myosin heads with a spacing d (~14.5 nm) located on opposite sides of the M-line. Since the polar arrays are separated by a bare zone of length *B* (~160 nm), the centers of pairs of myosin head layers on opposite sides of the M-line are separated by a distance of *L = B* + 48*d*. If *f* is an array of 49 myosin heads, and *g* is a set of two points (the centers of the arrays), then the structure is represented by the convolution *f ⊗ g*. The Fourier transform of a convolution is the product *F* (*f*)*G*(*g*) of individual Fourier transforms, which may have a minimum in the middle, depending on the relationship between the interference distance *L* and the myosin spacing d. If *L* is exactly equal to a half-integer multiple of d, then the reflection will be split into two peaks of equal intensity. The Fourier transform *F(f*) has a strong peak at the reciprocal spacing 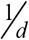, which is around 1/14.5 nm^-1^ (for M3), while *G(g*) is a fast-oscillating function (sin x/x)^2^ with a period 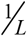. The fact that the doublet of two comparable intensities is observed suggests that *L* must be close to the half-integer multiple of d (Juanhuix *et al*., 2001; Linari *et al*., 2000).

The ratio of split peak intensities of the meridional M3 reflection is a very sensitive function of the interference distance L, implying that small changes in the axial motions of attached myosin heads (Linari *et al*., 2000) can be measured in response to filament sliding. This sensitivity was exploited in time-resolved X-ray interference experiments (Piazzesi *et al*., 2002; Huxley et al., 2006), aimed at measuring the motions of myosin heads during recovery from a quick release. The investigators applied a set of quick release steps to the muscle fibers, and for each of the steps, measured the positions and intensities of the higher-and lower-angle peaks of the M3 reflection. Separate measurements were performed before the release, at 110 μs following the release (which corresponded to the T_1_ phase) and at 2 ms following the release (T_2_ phase). The initial sarcomere length was 2.13 μm, which is in the range of maximal filament overlap. The data of Piazzesi et al. (2002) are reproduced in (Figure 1) in the form of two plots, the first one showing the ratio of intensities (I_HA_/I_LA_) of higher-and lower-angle peaks of the M3 reflection versus the length of release steps, while the second plot shows the spacings (S_HA_ and S_LA_) of the peaks versus applied length step. These data enabled Piazzesi *et al.* (2002) to calculate (with 0.1 nm precision) the average axial motions of the centers-of-mass of myosin heads (Δc) in the T_1_ and T_2_ phases as a function of release step distance. The centers of mass of the myosin layers on either side of the M-line do not significantly move between the T_1_ and T_2_ stages (see the discussion by these authors of their figure 3a), ruling out the rapid attachment/detachment hypothesis (see Figure 1).

**FIGURE 1.**
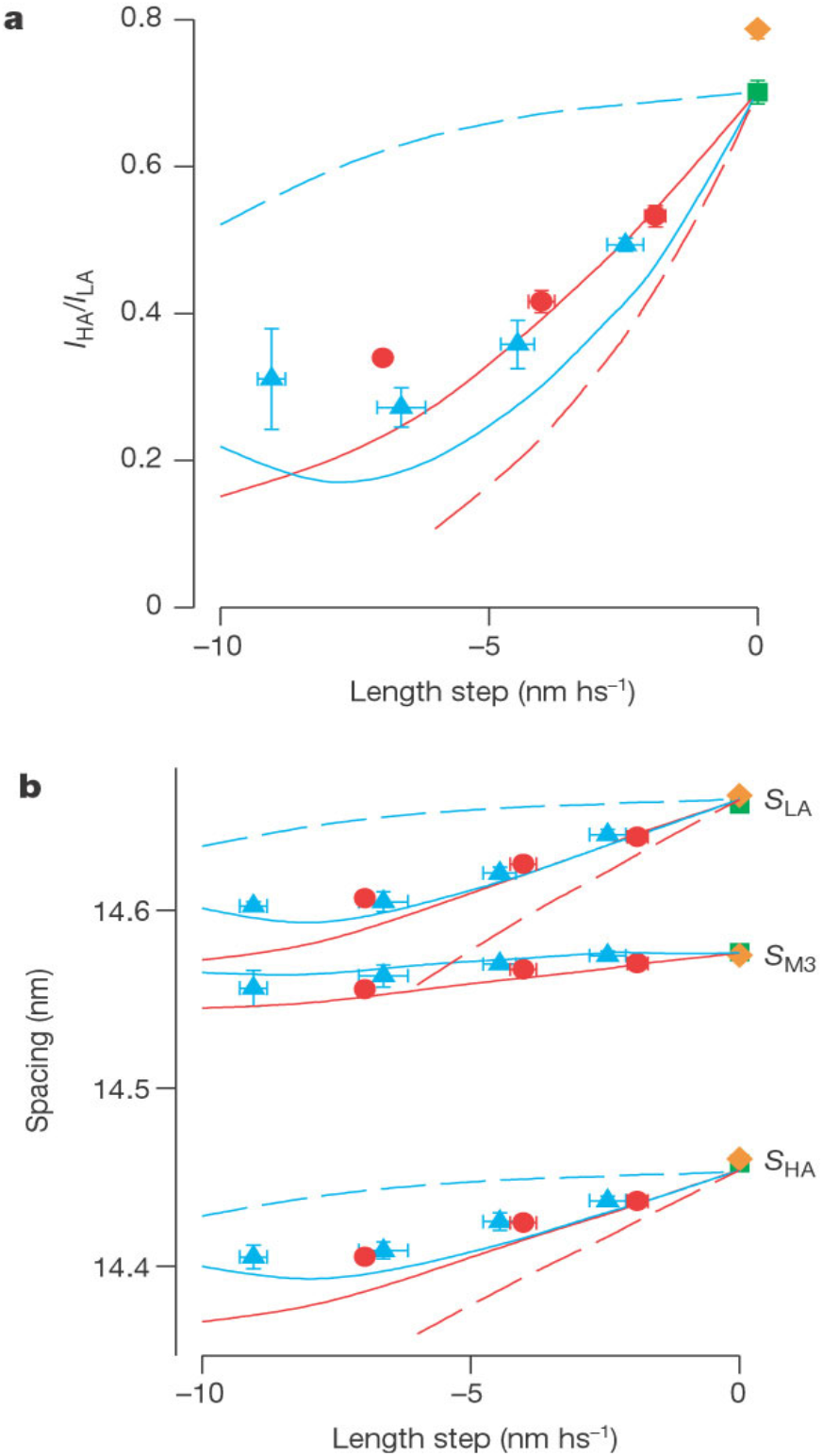
X-ray interference measurements on contracting muscle fibers during a quick release (Piazzesi et al, 2002). a. Intensity ratio of the higher and lower angle peaks of the M3 reflection plotted as a function of length step per half-sarcomere for the T_1_ (filled red circles) and T_2_ (filled turquoise triangles) phases of tension recovery. Best fits for the ‘one-headed’ swinging crossbridge theory (T_1_, solid red line) and the ‘two-headed’ version (T_1_, dashed red line). The best fit for the multiple myosin power-strokes theory (dashed blue lines) is well outside the measured data. b. X-ray spacings for the M3 reflection as a function of length step per half sarcomere. S_LA_ and S_HA_ are the spacings of the lower and higher angle peaks respectively, and I_M3_ is the un-split peak position. Color coding is the same as in 1a. Isometric, T_0_ pre-release, value (gold-filled diamond). (Permission to reproduce figure granted by Dr. Malcolm Irving).

At the time of the original experiments by Huxley and Simmons (1971), it was assumed that actin and myosin filaments were virtually inextensible, implying that myosin cross-bridge displacements would be equal to the applied length change or, equivalently, the relative amount of filament sliding. The experimental data of Piazzesi *et al.* (2002), however, show convincingly that the centers of mass of the myosin heads only shift by about a fifth of the applied length step (Figure 1). When more recent estimates for filament elasticity are used, the cross-bridge theory still leads to calculated displacements of myosin heads significantly (more than by a factor of 2) larger than observed by x-ray interference.. Such a serious discrepancy was true for both T_1_ and T_2_ stages and could not be eliminated when the elastic constants of the filaments were varied widely (Piazzesi *et al*., 2002). A recent model by Knupp et al (2019) allows for a very high actin-bound myosin head compliance (4.0 pN/nm) as long as the unbound heads take up strain, along with the thin and thick filaments.

Piazzesi et al. (2002) hypothesized that perhaps only half of the myosin heads in the overlap region respond to a length step, as later assumed by Knupp et al (2019), effectively multiplying the theoretically predicted values of the average axial motions of force-producing myosin heads by a factor of 0.5. Their justification comes from the fact that each myosin molecule has two heads, and while one of them can be attached to actin and shift during the quick release, the other one might be detached and not respond to the shortening step. Yet, since the two heads are joined at the head-tail junction, it was suggested that the paired, but detached head, might yet possess sufficient axial order to contribute significantly to the M3 reflection. This assumption appeared to reconcile the swinging cross-bridge theory with the calculations based on later time-resolved x-ray interference studies on muscle activation (Reconditi et al., 2011), where it is estimated that about 30% of the myosin heads are attached. However, the splitting of the M6 x-ray reflection could not be easily explained (see below), and the time dependence of changes in the positions of troponin and tropomyosin following Ca++ activation were reckoned not to be in agreement with prevailing theories.

Even given the assumption of coherent scattering by the unattached paired head, the fit to the data presented in the quick release experiments of Piazzesi et al. (2002) is not good at higher values (greater than 0.7 nm) of the quick release distance for the T_1_ phase (Figure 1a; reproduced from fig 2a, Piazzesi et al., 2002) and the modeled curve falls systematically below the experimental points for the T_2_ phase, suggesting that the myosin head displacements are too small to be accounted for by the swinging crossbridge theory.

**FIGURE 2.**
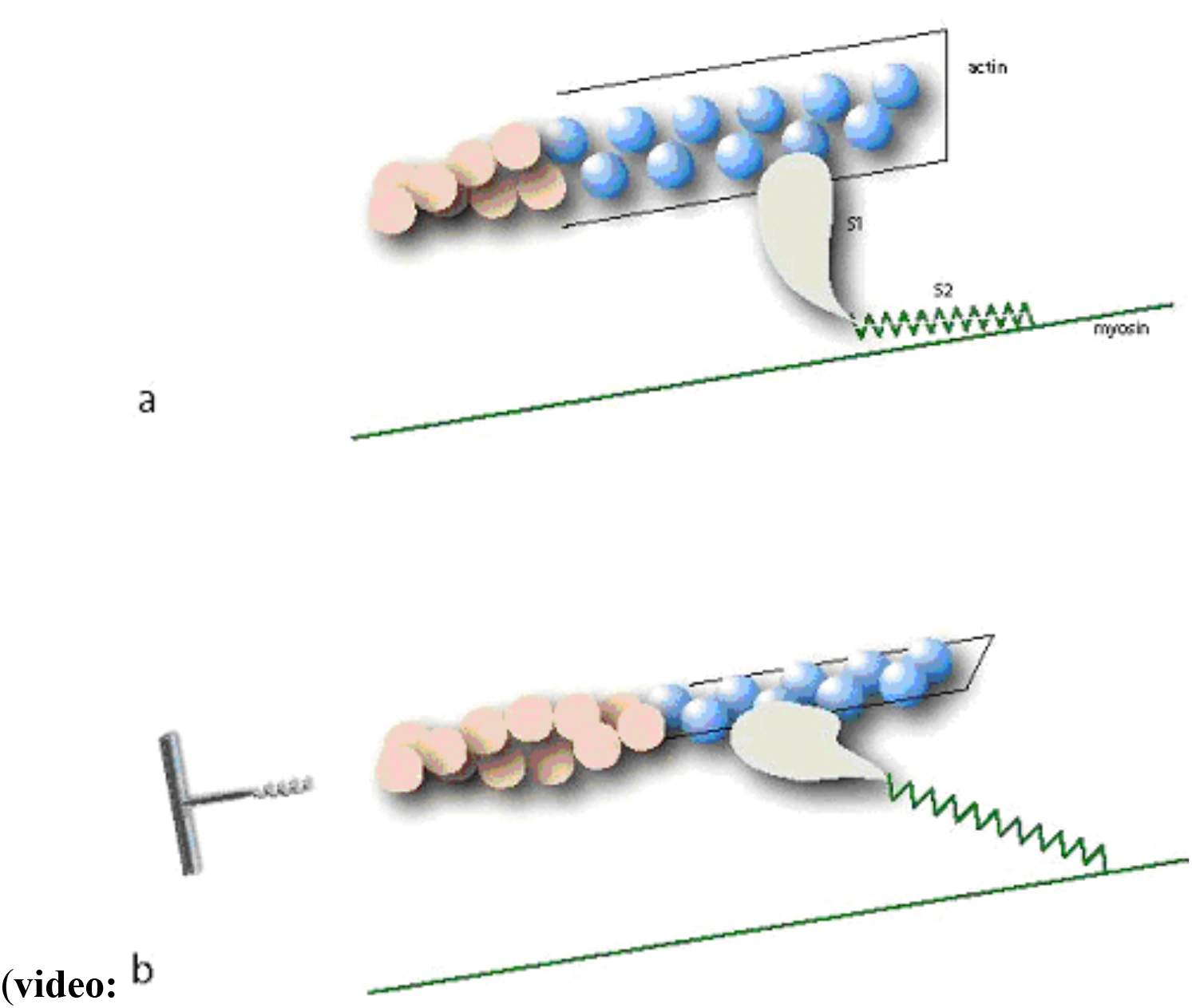
Cartoon exaggerating the effect of ribbon-to-helix transitions on azimuthal rotations of actin-bound myosin heads about the thick filament (torsional coupling effect). Grey: Myosin S1-domains; green: myosin S2-links; blue, actin ribbon subunits (r-actin); salmon, helical actin (h-actin). As r-actin subunits twist into the h-actin state bound myosin S1-heads rotate with ribbon segments. 2a. S1-head in a relatively unstrained state. 2b. S1-head twisted by approximately 60 degrees just prior to being twisted off (not to scale).

The spacings of the M3 (S_M3_) and M6 (S_M6_) reflections (Huxley et al., 2006) both increase by 1.5% during the development of a contraction, much higher than expected given literature values for the filament compliances and follow different time courses. Furthermore, the intensity of the M6 is much less sensitive to quick releases and other changes that affect myosin head form factors than the M3 (Reconditi et al., 2004; Huxley et al., 2006). This is surprising considering that it is the second order of the primary reflection (~14.3 nm) from successive layers of myosin heads along the thick filament. Indeed, Huxley et al. (2006) concluded that the ‘… M6 mainly arises from another component of the myosin filament, and that the myosin heads make only a small contribution to this reflection’. Since S_M6_ is a strictly linear function of the force independently of how muscle fibers are perturbed, whether by temperature jumps, applied length changes, or tension drops (Piazzesi et al., 2002; Reconditi et al., 2004; Linari et al., 2005), whatever the postulated structure on the thick filament contributing to the M6 might be, its role in force development needs to be explicated.

Huxley et al. (2006) were unable to find any actin-bound cross-bridge positions *including deployments of the unbound (paired) heads* that could explain why the ratio of the lower angle peak intensity (I_M6_) was always higher than the higher angle intensity (I_M6_), even considering models that accounted well for the fine structure of M3. Notably, electron density maps calculated using the four peaks of the M3 and M6 reflections showed an ‘extra density’ on the bound heads at higher radius (Knupp & Squire, 2009; Piazzesi et al, 2011) consistent with the reversal of I_HA_/I_LA_ for the M6 reflection. It is proposed below that the density at high radius can be attributed to bound r-actin segments having 14.3 nm longitudinal order.

### Actin-based model of muscle contraction

We examine here x-ray interference experiments carried out with protocols based on the seminal Huxley-Simmons (1971) measurements. In doing so, consideration will be given to the compliances of all of the components of the system.

During a quick release, the decrease in sarcomere length is accompanied by a relative translation of thin and thick filaments. In the cross-bridge theory, filament sliding is necessarily associated with axial motions of myosin heads, which bind and push actin filaments towards the center of the sarcomere. However, certain new effects become possible once structural transitions in actin are considered. One of them is a mode of filament translocation that can occur, in principle, *with very little axial movement of myosin heads.* Ribbon segments (r-actin) shorten to helices on the Z-line side of a bound myosin head (helicalization) while helical actin (h-actin) extends to r-actin on the M-line side (ribbonization) successively subunit-by-subunit as thin filaments translocate towards the center of the sarcomere. For each r-actin monomer that shortens on the Z-line side an h-actin monomer on the M-line side of the myosin head stretches to an r-actin subunit due to the binding of a myosin head further down the sarcomere (Schutt & Lindberg, 1993; Schutt & Lindberg, 1998). Because of the difference in length between the two structural forms of actin (Δx_r→h_ = 0.83 nm per monomer), an actin filament on the Z-line side of a bound myosin head shortens but because it extends by an equal amount on the M-line side there is no resulting net change in filament length. It should be noted that in the original publication of this model (Schutt and Lindberg, 1992) actin filaments in the resting state were depicted (figure 1) as being completely in the ribbon-state, the purpose being to illustrate commensurability of actin ribbons with the myosin lattice. X-ray measurements on the 5.9 nm actin-based reflection (Matsubara et al., 1984) and intensity increases in the first actin layer-line occurring on the same time scale as decreases in the ‘myosin’ layer lines (Huxley et al, 1983) were taken to be consistent with the hypothesis that some ribbon-to-helix transitions occurred during activation but the extent was not known.

Tension developed by this mode of filament sliding (“crawling”) is generated by actin itself. A pictorial example is shown in (Figure 2 and in accompanying video), where each consecutive “snapshot” represents the propagation of helicalization and ribbonization waves by exactly two monomers, which corresponds to the filament sliding distance of 1.66 nm (2*0.83 nm). The actin monomer to which the myosin head is bound is highlighted. The myosin head stays attached to an actin monomer until it is ‘twisted off’ by the rotation of a ribbon segment or when reached by a helicalization front. Thus, in this mode of filament ‘sliding’, attached myosin heads do not themselves rotate to generate force, but instead display small axial (Piazzesi et al, 2002) and large azimuthal (Huxley et al., 1983; Reconditi et al, 2004) passive responses to the actin force generators. Detached myosin heads can undergo conformational changes during ATP bonding and hydrolysis as they reset themselves for a further round of activating transitions in actin. The twisting off of myosin heads from actin is best seen in the accompanying video:

VIDEO: this video is frozen in the .pdf version. See Supplemental File No.1 for moving image)

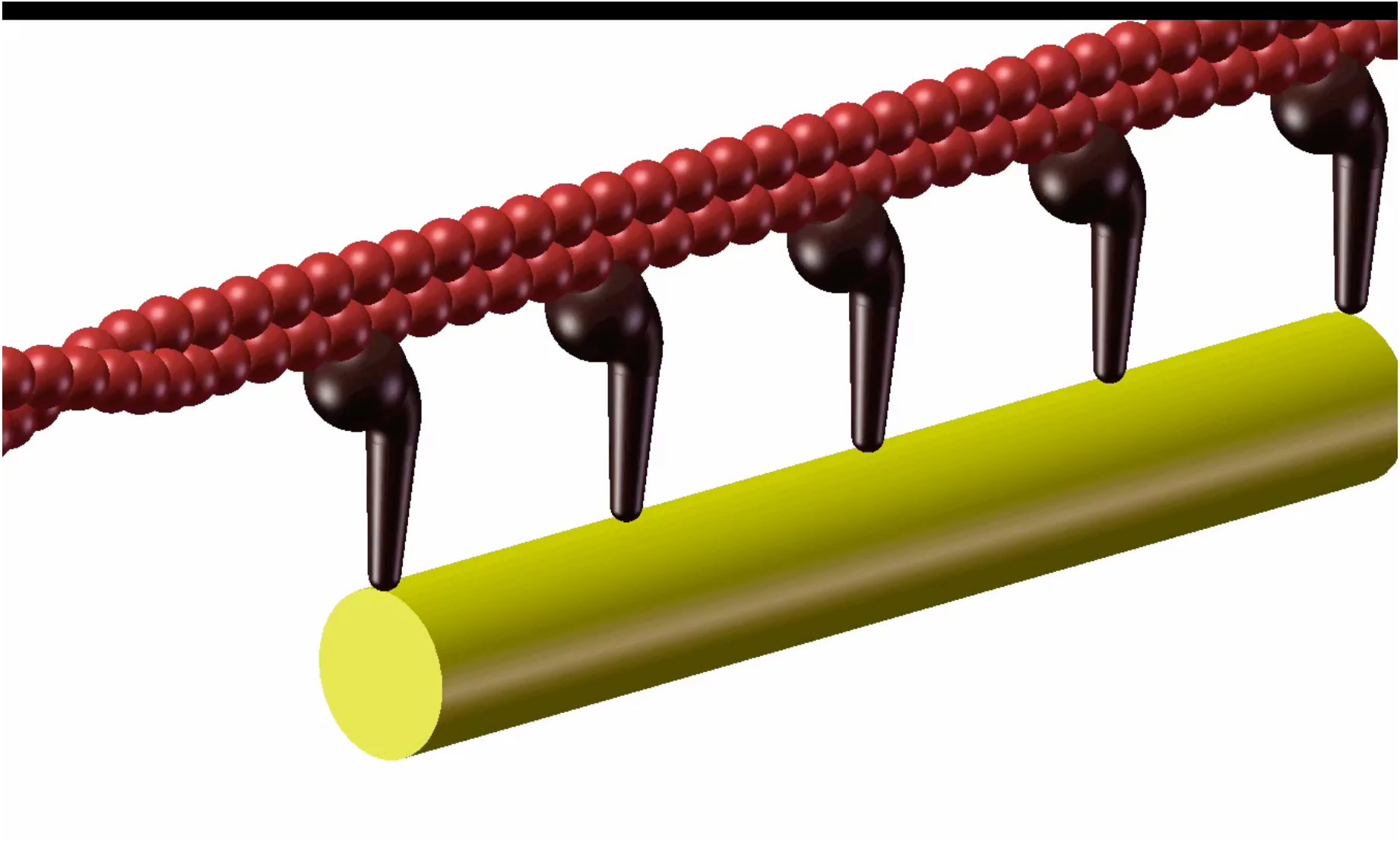

The proposed biochemical cycle of the actin power-stroke theory (Schutt *et al*., 1995a; Schutt & Lindberg, 1998) incorporates as an essential feature the ATPase activity of actin. The cycle starts with a myosin-ADP-Pi cross-bridge binding to an h-actin monomer (Lymn & Taylor, 1971), relieving the binding to tropomyosin. This leads to phosphate release from myosin and to the cooperative transformation of adjacent actin monomers into the metastable ribbon state, accompanied by ATP binding and hydrolysis to ADP:Pi. successively on each subunit of actin. The reverse of this reaction, the r-actin to h-actin transition, generates local forces of 100 pN per subunit as phosphate is released and h-actin rebinds tropomyosin. This model incorporates both a myosin ATPase (to activate transitions in actin) and an actin ATPase (to produce mechanical work).

Notice that the model on Figure 2 pertains only to the T_1_ phase, in which sarcomeres are shortening. In the T_2_ phase, in which sarcomere length is clamped and actin filaments cannot crawl, tension redevelops by a ‘churning’ action in which the axial shortening in actin subunits is counteracted by the torsional resistance of ribbon-bound myosin heads. Since the Z-disc is held fixed in an isometric contraction, *myosin heads would theoretically be pulled back towards the Z-line* by shortening actin subunits if it were not for the rotation of planar ribbon structures ‘downstream’ that twist bound myosin heads. The axial component of this twist pulls myosin heads in the opposite direction (towards the M-line). Tension builds as more and more strain energy is stored in the azimuthally strained (shear stressed) myosin heads, even though actin filaments are prevented from moving towards the M-line. Thus, the translation and rotation in actin subunits, coupled to the release of free energy from the actin ATPase, is used to produce the work done in distorting attached myosin heads and provides the mechanistic basis for understanding why there is no net change in the axial positions (Δc) of myosin heads between T_1_ and T_2_, as observed by Piazzesi et al (2002) and confirmed by our calculations (see ‘Calculations and Results’).

Thus, axial motions in myosin cross-bridges are expected to be smaller with an actin-based theory, and thus in better conformance with experiment, as compared to the swinging cross-bridge theory because, firstly, shortening within actin filaments themselves during the release takes up a significant fraction of the slack during the T_1_ phase of the recovery; and, secondly, in the T_2_ phase of recovery, where actin filaments are restrained from moving, the coupled force-generating transitions of translation and rotation result in little change in the axial positions of myosin heads. As will be shown below, the reduced axial bending in myosin cross-bridges during recovery from a quick release results in a much better fit to the x-ray interference data (i.e the curves of I_HA_/I_LA_ versus applied length step) of Piazzesi et al. (2002), especially for longer releases, compared with the cross-bridge theory.

### Modeling structural transitions in actin

Structural transitions in actin proceed simultaneously with the elastic response of filaments and cross-bridges to the quick release. Therefore, the change in half-sarcomere length (S) during a release is given as a sum of two components:

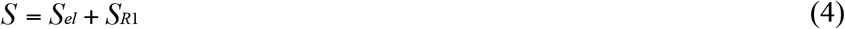

where S_el_ is the elastic component of half-sarcomere shortening, and S_R1_ is that due to the shortening of actin filaments themselves. In the cross-bridge theory, S_R1_=0. Similarly, the length by which filaments slide due to structural transitions in actin during the T_2_ phase will be denoted as S_R2_; S_R2_=0 in the crossbridge theory.

A.F. Huxley and colleagues (Ford *et al*., 1981) have derived expressions for the elastic response of the filament/cross-bridge system to changes in tension (see Supplementary Material). Considering that the elastic extension of a half-sarcomere is equal to a product of its compliance and the change in tension, and disregarding Z-line compliance, we see that the elastic change of the half-sarcomere length is a sum of contributions from strains in actin filaments, myosin filaments, and myosin cross-bridges in response to a change in tension:

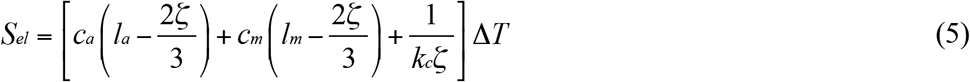

where ζ is the length of overlap zone, l_a_ and l_m_ – lengths of actin and myosin filaments respectively (within a half-sarcomere), c_a_ and c_m_ – actin and myosin filament compliances per unit length, k_c_-myosin head stiffness per unit length.

The ribbon-to-helix shortening contribution to sarcomere shortening, S_R1_, is the length by which filaments slide due to the contraction of r-actin during the quick release (see Fig. 2), and is equal to n*Δx_r→h_, where n is the number of r-actin monomers that are converted to h-actin. At higher loads these contractions will take place at multiple points along the thin filament. However, compensating transitions to the ribbon state of an equal number of actin subunits ensure that these contractions are coordinated so that the forces imparted to tropomyosin sum (see Schutt & Lindberg, 1992). Since larger length steps require more significant decreases in external load, the velocity of structural transitions in actin is higher for longer releases (Schutt & Lindberg, 1998). Considering that S_R1_ is the product of the velocity of ribbon shortening and the time, it will increase with increasing length steps. Assuming that the relationship between S_R1_ and the applied length step is linear:

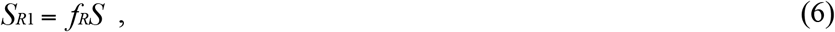

where the coefficient f_R_ is the fraction of half-sarcomere shortening attributable to ribbon-to-helix transitions, a free parameter to be determined (see Calculations & Results).

### Torsional resistance in myosin heads coupled to actin rotations

A crucial feature of this model is the rotation of actin filaments that takes place during contractions in actin subunits (Figure 2). Indeed, it is the cumulative rotation in the actin filament that stretches a bound myosin head about the thick filament axis, increasing the azimuthal strain in attached myosin heads. The force developed on bound myosin heads by ribbon segments can be resolved into two coupled components: an axial component, which pulls myosin heads towards the Z-line in the recovery phase, and an azimuthal component that rotates the plane of a ribbon segment around the filament axis, bending any attached myosin heads about the thick filament shaft (figure 2). The length S_R_ by which it shortens is related to the angle a by which the ribbon rotates:

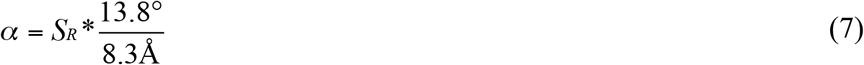

where, S_R_ is the generalized variable referring to the length by which a segment of r-actin contracts, irrespective of which phase of quick release is being discussed. If the T_1_ phase is being discussed specifically, then:

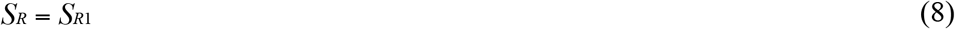

The total length of r-actin shortening from the beginning of the release to the end of the T_2_ phase will be the sum of individual shortenings during the T_1_ and T_2_ phases:

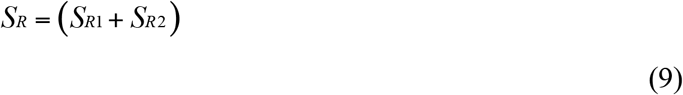

Ribbon segments respond to the torque developed by contracting actin monomers by rotating in the plane perpendicular to the filament axis. The axial component of the force generated by actin simply pulls myosin heads towards the Z-line, as some of the r-actin monomers located between each bound myosin head and the Z-line convert to the shorter h-actin state. We model the strain in myosin heads by a combination of rotation in the LCD converter domain (Dobbie et al., 1998) and a stretch of the S2-link. The tension borne by a cross-bridge through Hooke’s law is:

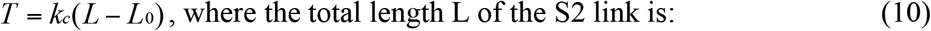

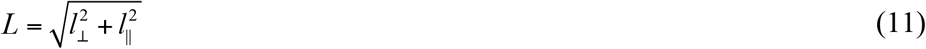

Since rotation is coupled to translation in this model, the extension of *l*_⊥_ is directly related to the increase in the length *l*_ǁ_ of the axial projection of the S2 link, which is given by:

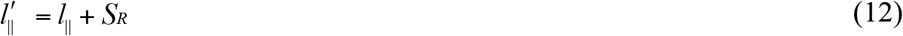

The stretching and bending of the myosin S2 link caused by the ribbon-to-helix transition during the T_2_ phase will reverse the shortening during the elastic (T_1_) response to the length step, restoring its length to its pre-release value. If bending occurs somewhere between the myosin catalytic and light-chain domains (Dobbie *et al*., 1998), a geometrical model (Figure 3) can be obtained that allows an expression to be derived for 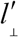 (assuming that the catalytic domain (S1) maintains a constant interface with actin), the modified value for the transverse projection *l*_⊥_ of the S2 link:

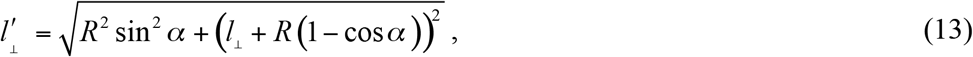

where R is the radius of actin filament plus the length of the myosin S1 catalytic domain, and a is the angle of the twist. As shown explicitly below (see Calculations and Results), this equation can be solved for each value of S to obtain an average value for S_R2_ (S).

**FIGURE 3.**
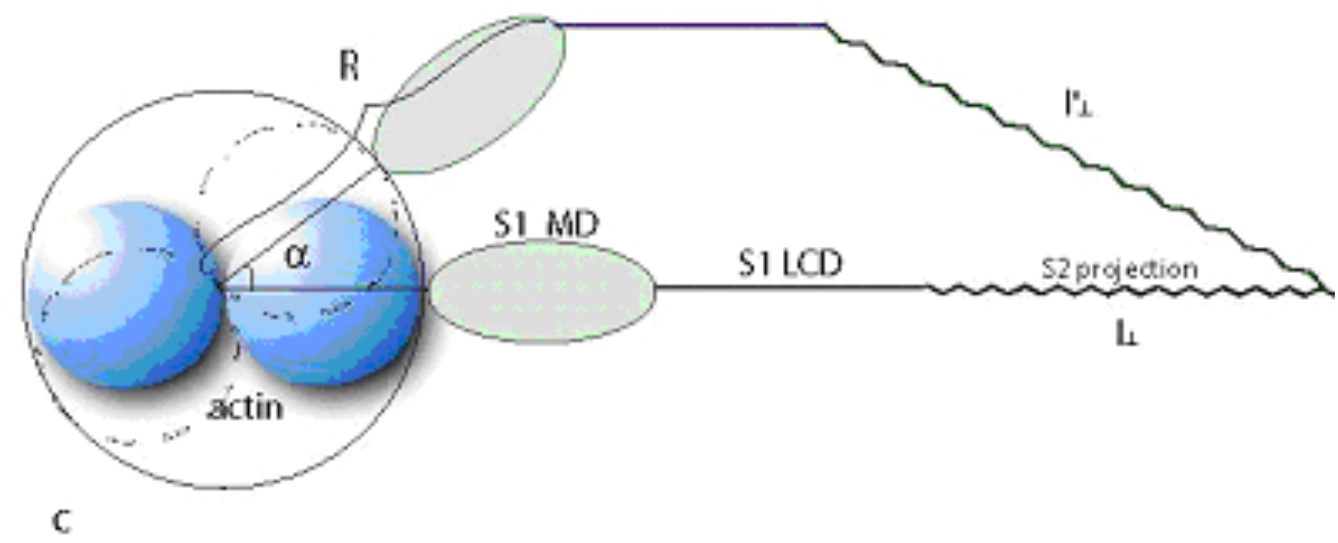
Model for apportioning strain in attached myosin heads. In the actin power-stroke model attached myosin heads are rotated about the axis of the thick filament as actin subunits twist in the ribbon-to-helix transition (torsional coupling). This results in the stretch of the S2 element of myosin, where 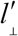 is the transverse projection of the S2 myosin link (see text). R is the combined radius of the actin filament (blue) and the myosin S1-head (stippled green). The angle α is the degree to which the ribbon is rotated under the action of contracting r-actin subunits. Not drawn to scale. Changes in the S1 light chain domain (S1-LCD) not depicted.

The azimuthal rotation of ribbon-bound myosin heads contributes to a stretching of the myosin elastic elements. In the absence of a detailed structure-based theory for strain accommodation in myosin heads, we treat the problem in terms of distortions in the myosin light chain domain (LCD) with a stretched tS2-link reminiscent of the original model of Huxley & Simmons (1971). It has recently been shown that the S2-link itself has a very low compliance (Brunello et al., 2014) so that our use of a stretched S2-link in calculating torsional resistance should not be taken literally but rather as an application of the virtual work concept showing that rotational distortions (+/− 17°) of the magnitude inferred from x-ray measurements on contracting muscle fibers (Huxley et al., 1983; Reconditi et al., 2004) provide a reasonable explanation for torsional resistance in bound myosin heads in response to ribbon-to-helix transitions. Future work will be directed at using crystal structures of myosin to calculate the strain energy in the LCD converter domain in response to actin-induced twists.

The movement of the myosin S2 link away from thick filament induced by the ribbon-to-helix transition during the T_2_ phase reverses geometrical changes in S2 during the T_1_ phase, assuming that the tension per crossbridge in T_2_ is the same as in the isometric state. Substituting S_R_ from (9) and taking (7) into account, we can obtain a non-linear equation for S_R2_, which is the length by which r-actin contracts during the T_2_ phase. By solving this equation for each value of S, we can obtain the function S_R2_(S), which is the response of the force-generating element in r-actin to the applied length step.

The S_R2_ (S) relationship is quasi-linear for small to moderate releases, reflecting the fact that contractions in actin take up about 30% of the applied length change (see below ‘The effect of actin shortening on length – tension curves’); however, for large releases (>5.0nm), shortening of r-actin during tension recovery does not increase proportionally to length step. Instead, as the release steps increase, tension regeneration in the T_2_ phase is limited, to a larger and larger extent, by the resistance to the torque developing in attached myosin heads as other ribbon segments in the fiber take up the slack. Furthermore, a larger fraction of cross-bridge connections are broken as tension recovers from large step releases. In terms of the motions of myosin heads between T_1_ and T_2_ phases, this model substitutes the large axial motions predicted on the basis of rotating cross-bridges by a combination of moderate axial and specific azimuthal motions. *Therefore, since the fine structure of the M3 myosin x-ray reflection depends only on axial motions of myosin heads, changes in M3 between the T_1_ and T_2_ phases are reduced to the extent that tension is stored azimuthally in the elastic elements of the myosin heads.*

### Dependence of myosin head detachment on transitions in actin

Stiffness in muscle fibers during recovery from a quick release is smaller than during an isometric contraction (Cecchi *et al*., 1982; Ford *et al*., 1985) indicating that some myosin heads detach during the quick release and ensuing tension recovery. The tension drop can be explained on the basis of the actin power-stroke theory since a myosin head detaches whenever passed by a helicalization front (Figure 4). Thus, a certain fraction of the heads will detach during both T_1_ and T_2_ phases of the quick release, and the probability of detachment generally is:

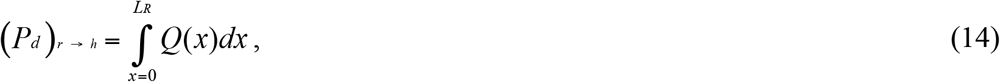

where L_R_ is the distance traveled by the helicalization front, and *Q* (*x*)*dx* is the fraction of ribbon segments with outside boundary within a distance interval *x ↔ x* + *dx* from the myosin head attached to it. We can determine L_R_ more specifically by using the values of rise-per-subunit of h-actin helix (*h* = 2.74*nm*) and the length difference between h-actin and r-actin monomers (Δ*x_r → h_* = 0.83*nm*). For each actin monomer converted from ribbon to helix, the actin filament translocates by the amount Δ*x_r → h_* relative to the myosin filament, whereas the helicalization front actually travels a greater distance *h* + Δ*x_r → h_*, so that:

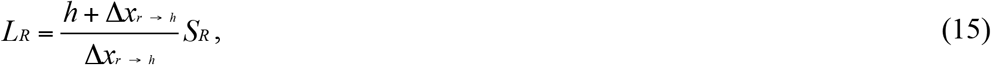

where S_R_, the length of filament translocation, is defined through (8) and (9).

**FIGURE 4.**
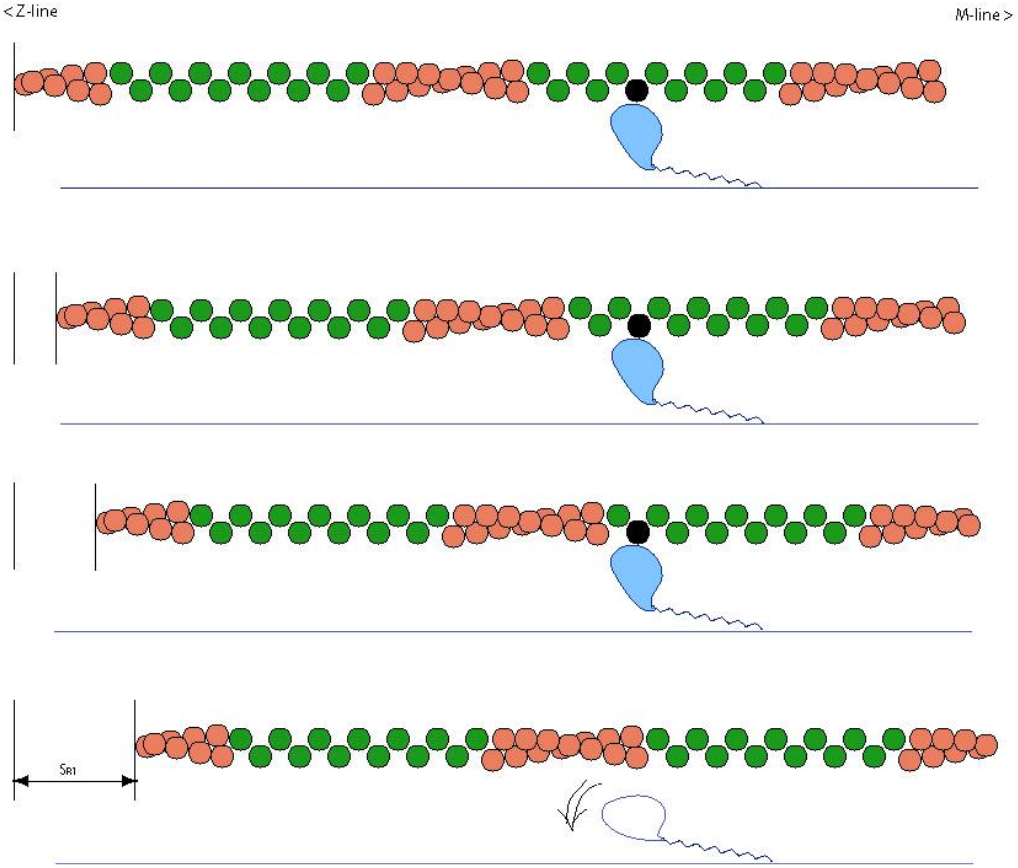
A cartoon, not drawn to scale, depicting myosin detachment from actin ribbon segments due to the ‘twisting off’ effect when a helicalization front (salmon) traveling along a crawling filament reaches the bound head (blue) from the left. See Schutt & Lindberg (1998) for a more detailed description. A myosin head (blue) bound to an actin subunit (black) in the top panel being released in the bottom panel when the helical wave passes through. This diagram does not show the rotation of ribbon segments (green) to the right of the advancing helicalization front as depicted in Figure 2 (torsional coupling). Both effects are present in sarcomeres during recovery from a quick release. **(video:** https://www.dropbox.com/sh/pmbowduod4gdz5f/AAAc1S6qa0GEstklkkySxBvra?dl=0)

A bound myosin head not reached by a helicalization front can either remain attached or can detach depending on the magnitude of the imposed twist associated with torsional coupling (Figure 2). In the latter case the twist is proportional to S_R_ through (7), so that the probability of detachment due to ‘twisting off is a function of S_R_:

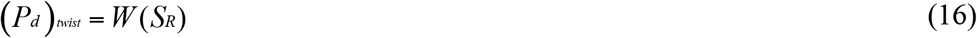

Combining (14) and (16) gives the probability of detachment as:

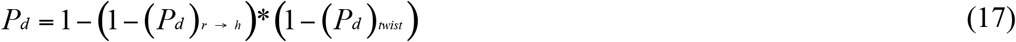

Re-attachment of those myosin heads that have detached during a quick release depends on the completion of another round of the myosin ATPase cycle (Lymn & Taylor, 1970), which is much slower than the T_2_ phase of tension recovery. It follows that the fraction of attached heads in the T_2_ phase will be smaller than in the initial isometric state. This reduction in the number of contributing force generators after a quick release implies that the recovered tension is proportional to the fraction of myosin heads that are still attached at the end of the T_2_ phase:

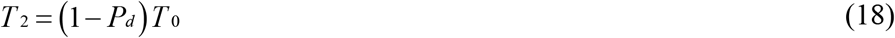

Notice that equation (18) assumes that tension borne by each attached cross-bridge is the same as during the T_2_ phase in the initial isometric contraction. In the present model, attachment of myosin heads becomes possible only in the later (T_3_ and T_4_) phases of tension recovery, during which tension fully returns to its isometric value.

The forms of the functions *Q*(*x*) and *W* (*S_R_*) have been left completely general, but it is appropriate to take functions with well-known switching properties such as:

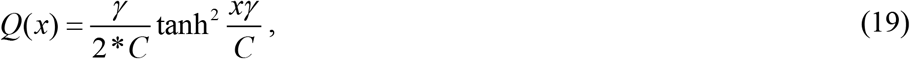

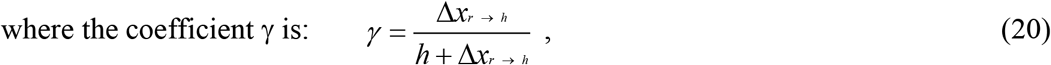

a function that integrates easily and gives

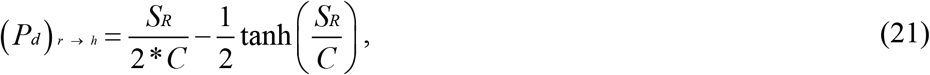

reminiscent of the original Huxley-Simmons (1971) analytical approximation of the T_2_ tension-length relationship (equation 3).

Since a ribbon segment is 429 Å long, it is certain that all myosin heads will detach when a helicalization front travels a distance 429 Å. This gives rise to the normalization condition for the distribution function *Q*(*x*):

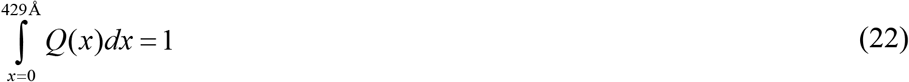

implying that C=3.33 nm.

As for *W*(*S_R_*), a smooth “switching” function suffices:

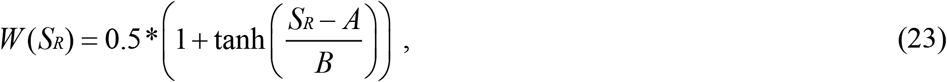

The parameters for the function *W*(*S_R_*) are A=3.33 nm and B = 0.83 nm.

*W*(*S_R_*) is the probability of myosin heads being twisted off by ribbon-to-helix transitions in actin. The switching function for *W*(*S_R_*) implies that heads tend to detach in a certain range of the twist angle. The parameter A is a characteristic length of shortening of r-actin, at which 50% of the heads would detach. Taking A=3.33 nm is equivalent to saying that 50% of the myosin heads would detach when 4 (3.33 nm/ 0.83 nm) actin monomers (i.e., one-third of a ribbon segment) have undergone transformation from ribbon to helix. The parameter B determines the half-width of the *W*(*S_R_*) relationship, where B= 0.83 nm entails an uncertainty of ±1 monomer, so that most (>80%) of the heads detach when the number of actin monomers already converted from ribbon to helix is between 3 and 5. Due to torsional coupling, the length of r-actin shortening is proportional to the twist in the actin filament, so that the characteristic twist at which myosin heads tend to detach is A*(13.8°/0.83 nm), which is around 55° when A=3.33 nm. The prediction that a ribbon-bound myosin head will detach with high probability at twist angle around 60° implies that a different myosin head on the opposite side of the ribbon projecting from a different myosin filament in the hexagonal lattice will be in position to bind with sharply-defined stereochemistry. The picture that emerges is that crawling actin filaments, centered on trigonal positions in the myosin thick filament lattice, regulate the binding of myosin heads by a synchronous mechanism that engages and releases myosin heads on neighboring thick filaments in succession one after the other, somewhat akin to a rotary combustion engine. Such a self-balancing system ensures that actin filaments are bound to myosin heads at virtually every moment during the isometric contraction, which is critical for the maintenance of tension.

Finally, although the functions *Q*(*x*) and *W*(*S_R_*) are similar in form, the processes they describe are fundamentally different. The distribution function *W*(*S_R_*) represents the probability of detachment of myosin heads as a function of r-actin shortening purely as a statistical effect depending on the cumulative rotation of the ribbon to which the head is bound. The function *Q*(*x*), on the other hand, is the distribution of helicalization fronts relative to bound myosin heads. Detachment corresponding to this effect is deterministic and occurs when a bound myosin head is reached by a helicalization front (Figure 4 and accompanying video). Although the exact forms for the functions *Q*(*x*) and *W*(*S_R_*) were chosen primarily for their mathematical properties., they represent well the qualitative expectations of the model, and allow an explanation of the T_2_ tension-length curve purely as an effect of myosin heads detaching in response to ribbon-to-helix transitions in actin. *Q*(*x*) is the distribution of helicalization fronts approaching actin-bound myosin heads from a certain distance x. We hypothesize that the nonlinear statistical distribution (19) is the result of cooperativity in structural transitions in actin. The argument x of *Q*(*x*) decreases from 42.9 nm to 0 nm as the helicalization front approaches a bound myosin head. The form of the distribution (19) means that, at a given moment of time, there is a bigger chance of finding ribbon segments that have just bound myosin heads, rather than ones that are about to twist off. Such a tendency again results from cooperativity in the actin ATPase cycle, which entails the velocity of propagation of structural transitions in actin is lowest when a ribbon segment has just bound a myosin head, and gradually speeds up as it shortens. Such an effect gives rise to the statistical decrease in the population of ribbon segments with small values of x.

## Results and Discussion

### The effect of actin shortening on length tension curves

During the T_2_ recovery phase following a quick release in tension actin ribbon subunits will shorten a distance of S_R2_ per half sarcomere to develop force (Equation 9). A calculation showing the amount of ribbon shortening per half-sarcomere as a function of release distance (see Appendix A) based on the actin power-stroke model is shown in Figure 5.

**FIGURE 5.**
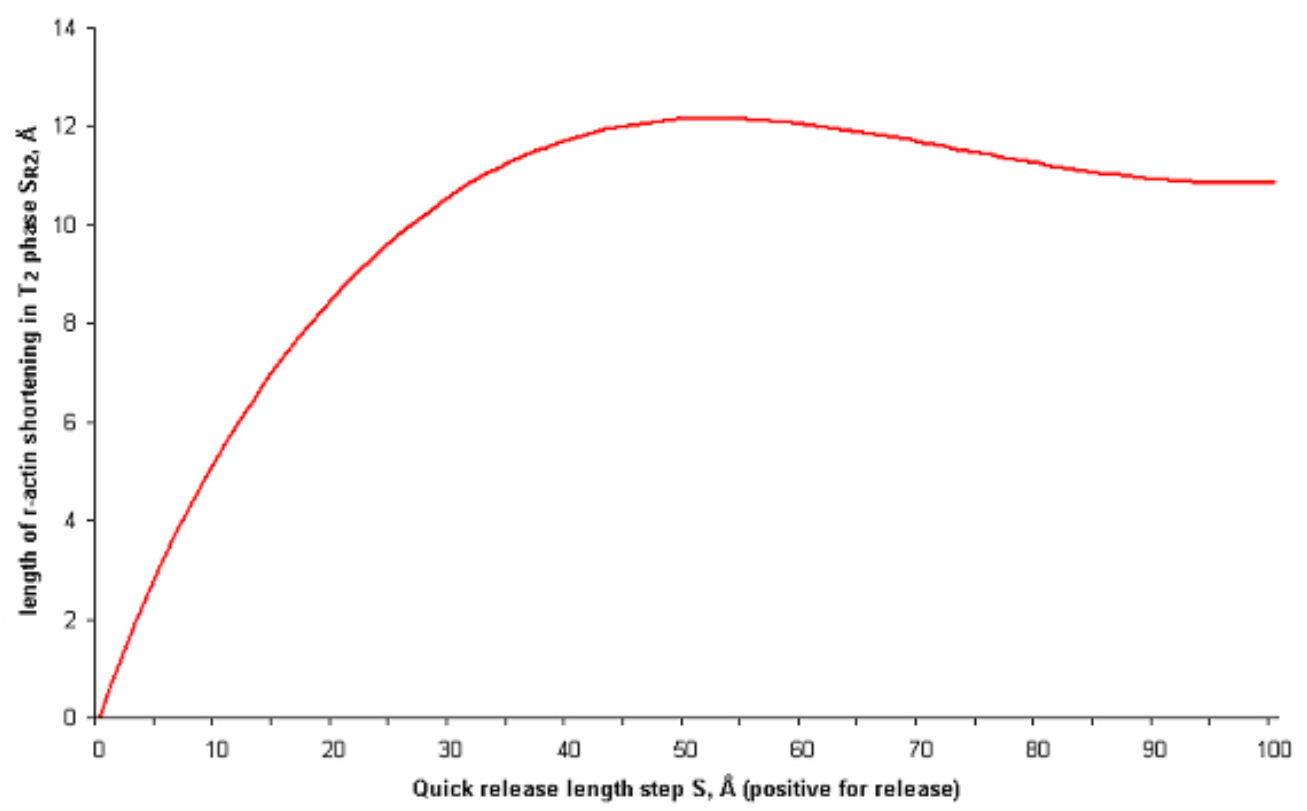
Total ribbon shortening (S_R2_) per half-sarcomere during the T_2_ phase of tension recovery as a function of length step (S).

### The T_2_ length-tension curve based on the actin power-stroke model

The direct comparison of our simulated T_2_ length-tension curve against experimental curves was complicated by the fact that experimental research groups use different methods for extracting T_2_ tension (Ford *et al*., 1977; Eisenberg *et al*., 1980). On (Figure 6) we compare our curve with the original analytical approximation (Huxley & Simmons, 1971). We can see that the shapes of both curves are very similar, although the intercepts on the length axis are different. In fact, analysis of more recent T_2_ tension-length curves (Chen & Brenner, 1993) suggests that the intercept is smaller than the one originally extracted by Huxley & Simmons and may be somewhat smaller than 10.0 nm.

**FIGURE 6.**
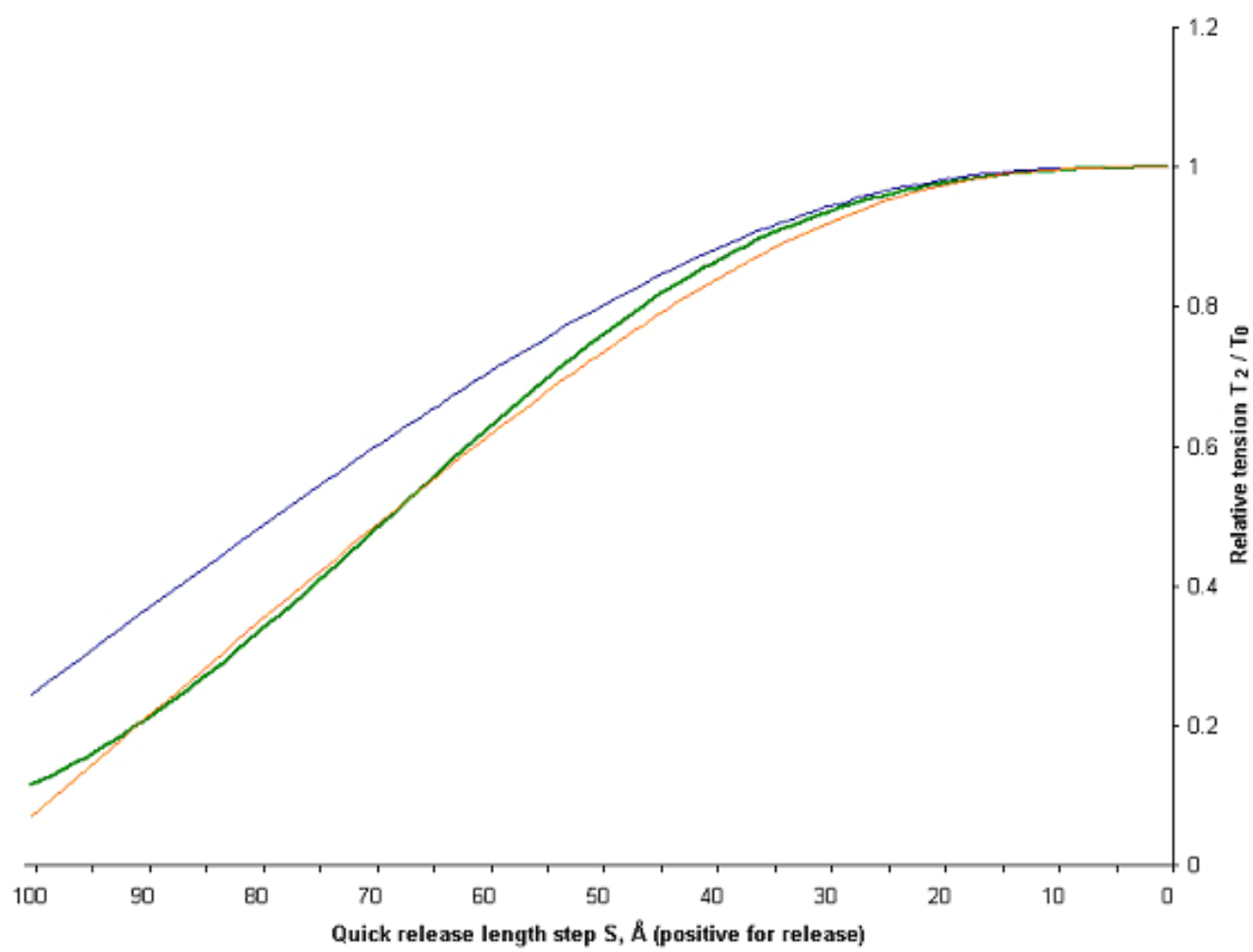
Length-tension curves for the T_2_ phase of recovery following a quick release. Calculated curve for the actin power-stroke theory (green); calculated with original Huxley-Simmons (1971) (green); Huxley-Simmons (1971) analytical approximation, with the following modification: parameter a increased from (2 nm)^-1^ to (1.75 nm)^-1^ to agree with later observations of smaller length axis intercepts (Chen & Brenner (1993).

### Modeling x-ray interference results

The formalism presented here can be used to model the recovery behavior of muscle fibers following the drop in tension caused by a rapid change in length. The calculated positions of actin-attached myosin heads can be used to model changes in x-ray interference diffraction patterns. The results of the calculations are summarized by plots of relative intensities and spacing of the higher- and lower-angle peaks of the M3 reflection versus length of the release step, which can be compared with data from the time-resolved X-ray experiments on muscle fibers (Piazzesi *et al*., 2002). The experimental and calculated ratios of higher- and lower-angle peaks of the meridional M3 reflection are shown in Fig. 7 (T_1_) and Fig. 8 (T_2_). The spacings of the higher- and lower-angle x-ray peaks are shown in Fig. 9. In the simulations that led to Fig.7 – Fig.9, the relative proportion of ribbon-to-helix contribution to sarcomere shortening was taken as f_R_=0.30; the disorder factor of the heads that detach during the quick release was estimated as 0.5; the elasticity of myosin heads was considered to be distributed between the S2 stretching mode and the light-chain domain tilting mode in a 1:1 ratio. In the calculation for the torsional coupling effect, we used the values of R = 100 Å, the initial (pre-release) length of the S2 myosin link L_0_ = 40.0 nm and the initial tilt of S2 link against the filament axis *φ* = arctan (0.28).

**FIGURE 7.**
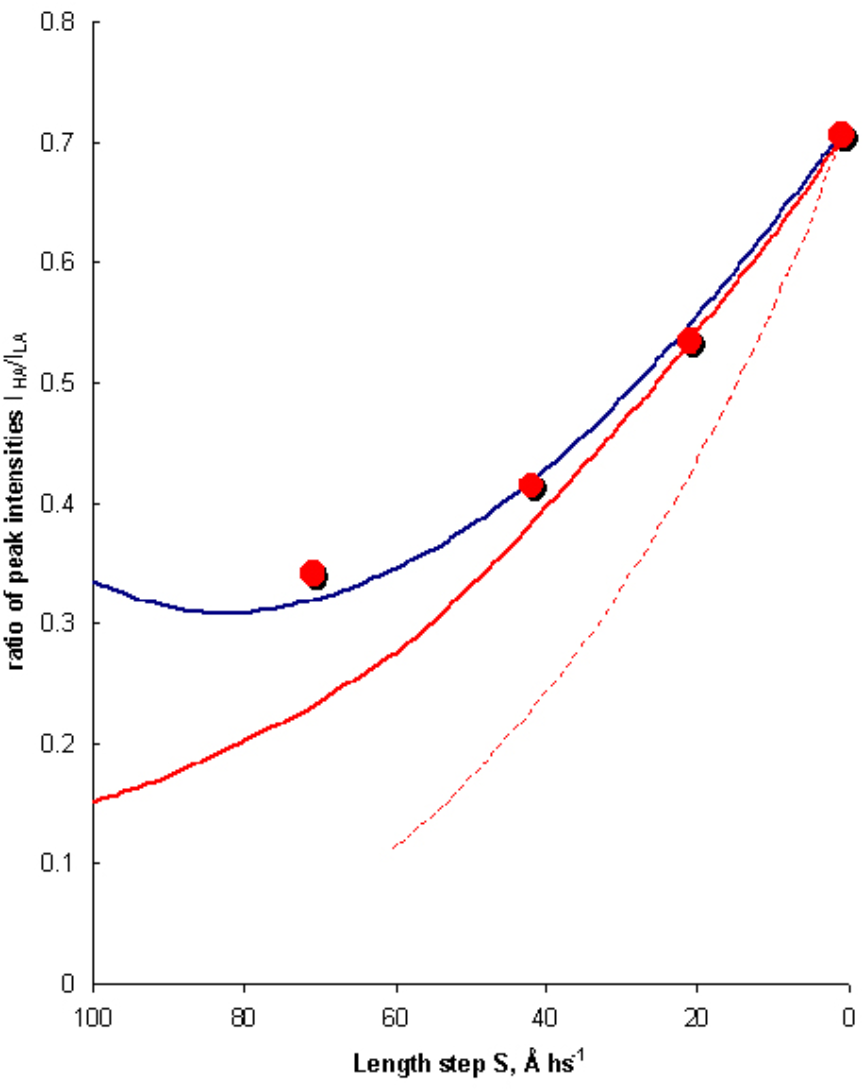
Comparison of model calculations with x-ray interference data of Piazzesi et al. (2002) during the T_1_ phase of tension recovery following a quick release. Ratio of the split peak intensities of the M3 meridional reflection (I_HA_/I_LA_) plotted as a function of step length per half-sarcomere (S). Measured values (Piazzesi et al., 2002) plotted as red circles. The best fit obtained under the assumptions of the cross-bridge theory with one head bound of the myosin dimer (red curve). The best fit obtained under the assumptions of the actin power-stroke theory (blue curve). The best fit with the rotating cross-bridge theory with both heads of the myosin dimer bound to actin (dotted red curve).

**FIGURE 8.**
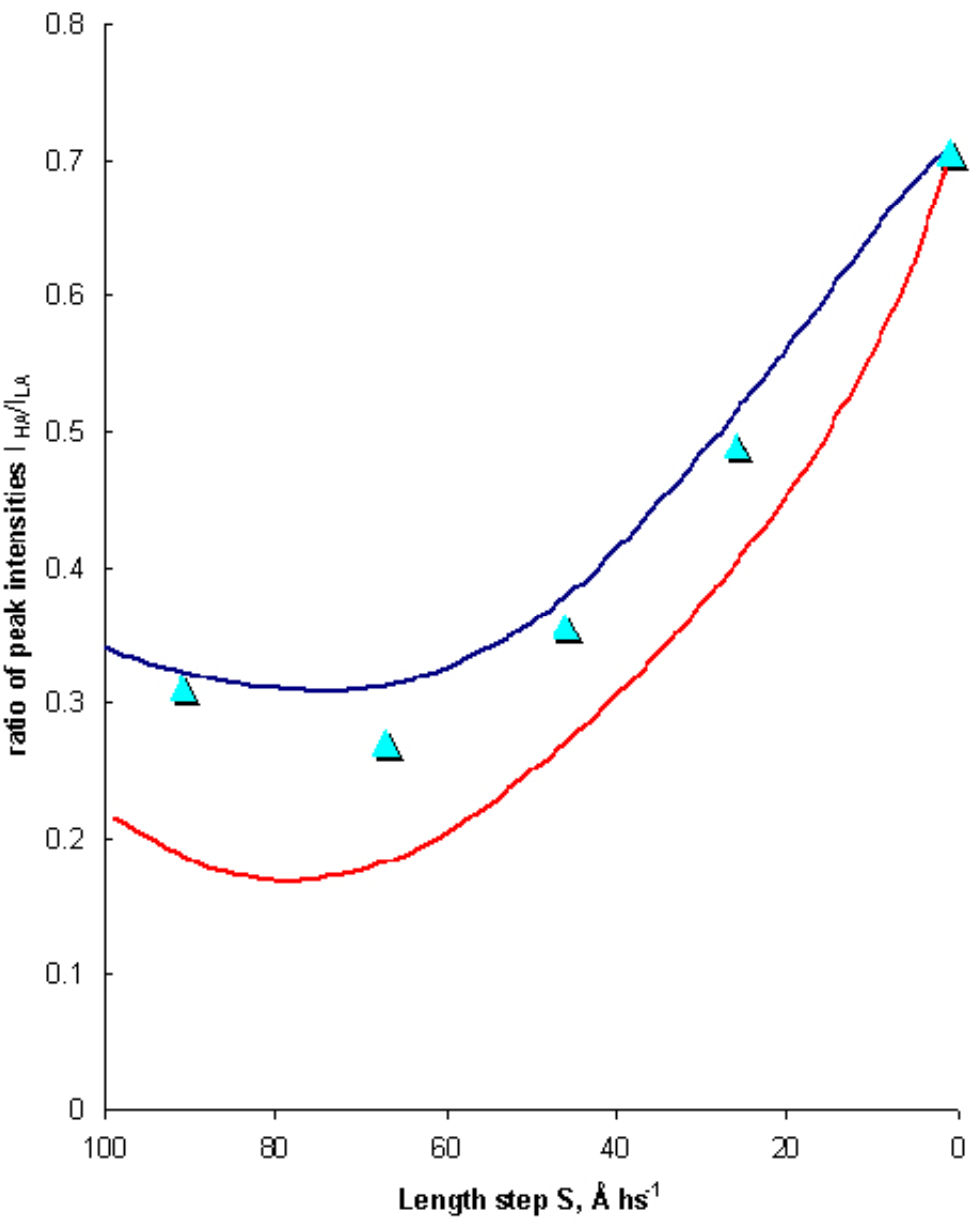
Comparison of model calculations with x-ray interference data of Piazzesi et al. during the T_2_ phase of tension recovery following a quick release. Ratio of the split peak intensities of the M3 meridional reflection (I_HA_/I_LA_)x plotted as a function step length per half-sarcomere (S). Measured values plotted as turquoise triangles (Piazzesi et al., 2002). The best fit obtained under the assumptions of the actin power-stroke theory (blue curve). The best fit under the assumptions of the rotating cross-bridge theory with one of the two heads of the myosin dimer bound to actin.

**FIGURE 9.**
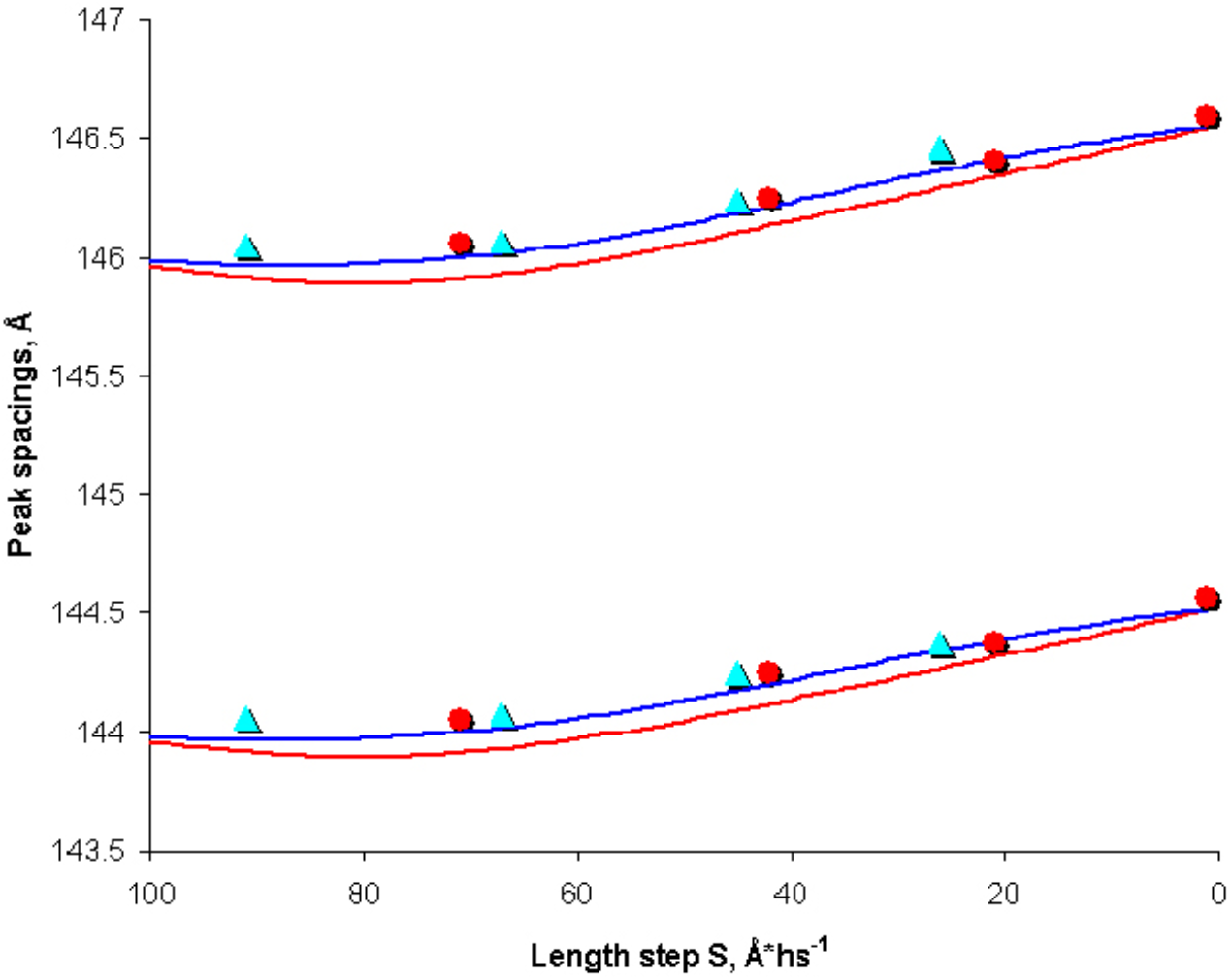
Peak X-ray spacings for the M3 reflection during the T_1_ and T_2_ phases of tension recovery following a quick releases as a function of length step. Red filled circles, experimental values of Piazzesi et al. (2002); calculated values based on the actin power-stroke theory (turquoise filled triangles). Red lines, calculated values based on the rotating cross-bridge theory (Piazzesi et al., 2002). Blue lines, best fit through calculated data points for the actin power-stroke theory.

### Choice of modeling parameters

The fits shown in Fig.7 – Fig.9 demonstrate that the actin-based model of quick release gives an excellent quantitative account for the x-ray interference measurements on contracting muscle fibers subjected to quick releases (Piazzesi et al. (2002)). The fitting of the experimental data required the setting of certain parameters, discussed below, including the proportion of shortening attributable to ribbon-to-helix transitions (f_R_), the disorder factor for scattering from detached myosin heads, the geometric parameters for calculating azimuthal actin-bound cross-bridge distortions arising in response to rotations in actin, and the contribution of actin ribbons to the M3 reflection. All calculations were performed over a range of choices for these parameters in order to optimize the fit to the experimental data and to assess to robustness of the model.

Fig. 10 and Fig. 11 show the effect of varying f_R_ on the relative intensity plots, for T_1_ and T_2_ respectively. It is clearly seen that the effect of the ribbon-to-helix contribution to sarcomere shortening displays itself in lifting both T_1_ and T_2_ curves up. The reason is that when larger fractions of sarcomere shortening are taken up by structural transitions in actin a smaller fraction is attributable to myosin head compliance. The resulting reduction in myosin head axial motions essentially shifts the ratio of peak intensities closer to its pre-release value. Fig. 10 and Fig. 11 show that the fit of experimental data visibly worsens when the values of fR move away from 0.30, for both the T_1_ and T_2_ curves.

**FIGURE 10.**
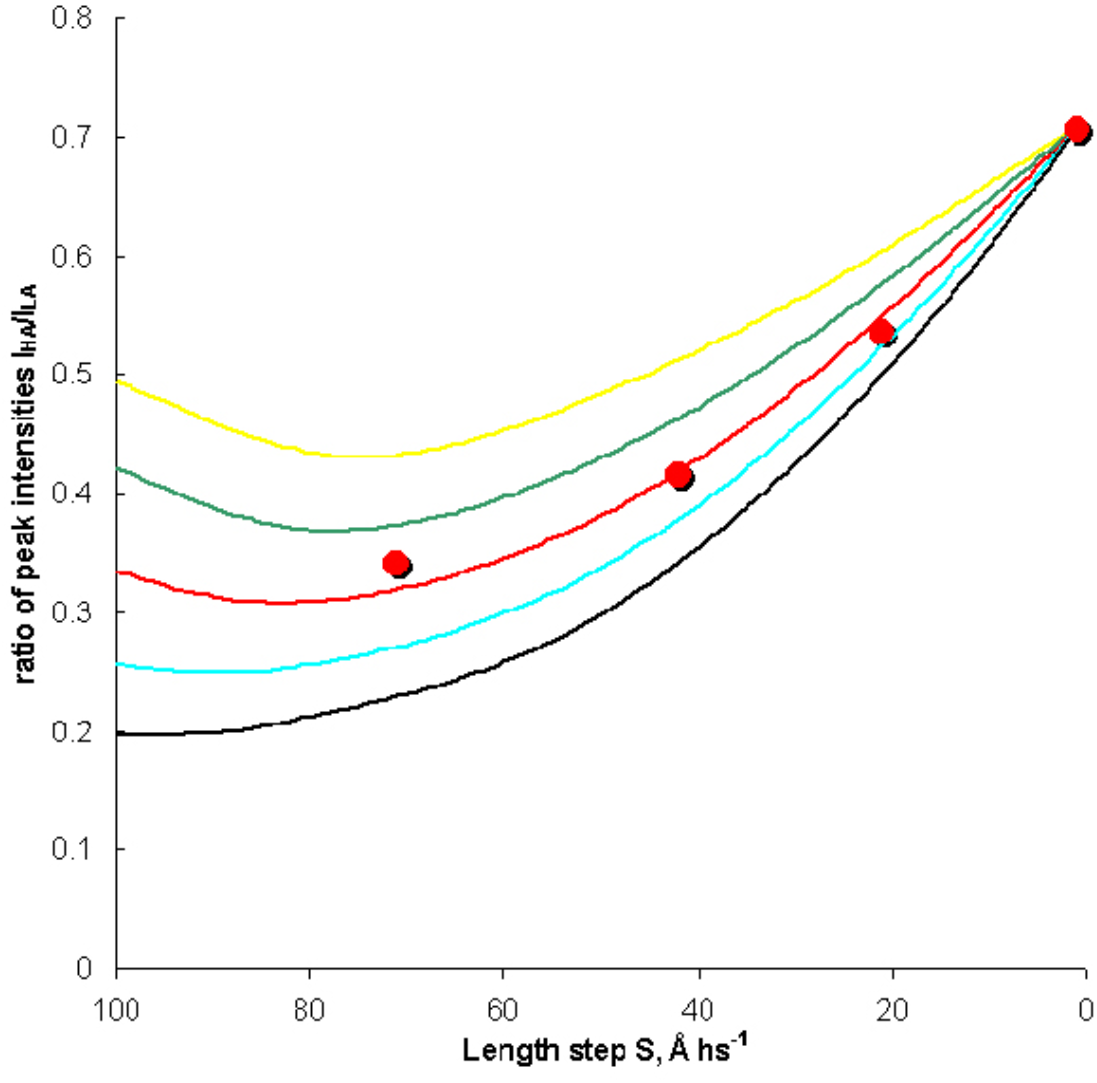
Sensitivity of fit to x-ray interference data (I_HA_/I_LA_) to choice of the fraction of sarcomere shortening attributable to ribbon-to-helix transitions (f_R_) during Ti phase of tension recovery following a change in step length. Measured values (Piazzesi et al., 2002) as filled red circles. The best fit is obtained with f_R_ = 0.3 (red). f_R_ = 0.4 (yellow), f_R_ = 0.35 (green), f_R_ = 0.25 (turquoise), f_R_ = 0.20 (black).

**FIGURE 11.**
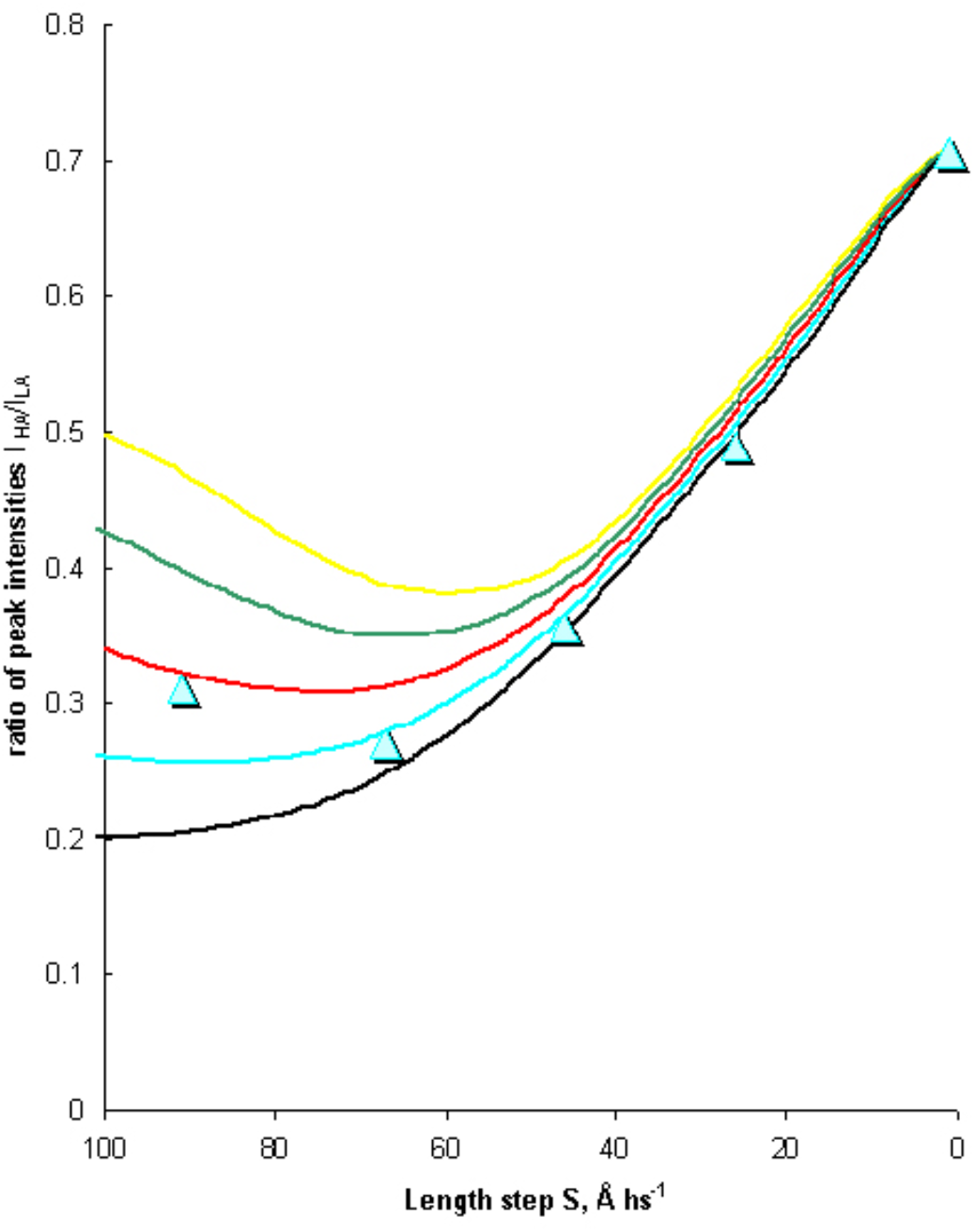
Sensitivity of fit to x-ray interference data (I_HA_/I_LA_) to choice of the fraction of ribbon shortening (f_R_) during T_2_ phase of tension recovery following a change in step length. Measured values (Piazzesi et al., 2002). The best fit is obtained with f_R_ = 0.3 (red), f_R_ = 0.4 (yellow), f_R_ = 0.35 (green), f_R_ = 0.25 (turquoise), f_R_ = 0.2 (black).

Similar sensitivity plots demonstrate the dependence of the fit to the x-ray interference data on the effective sum of the radius the actin filament together and the length of the attached myosin head (R, in figure 4), which is the variable used in calculating azimuthal motions in attached cross-bridges as they respond to torsions produced in force-producing ribbon-to-helix transitions (see Fig.4). As in the case of combined radius R, sensitivity plots (not shown) show that increases in the tilt of S2 links emphasizes the effect of torsions on the fit to the x-ray interference data. Our best value of the tilt (*ϕ* = arctan(0.28)) corresponds to the transverse projection of S2 links around 40*0.28 = 11.2nm. This estimate is reasonable, since the distance between the centers of thin and thick filaments is 24 nm (Huxley & Brown, 1967), which includes the radius of actin filament (~4 nm) and the sum of radial projections of both S1 head and S2 link. Estimating the radial projection of a myosin head as 10.0 nm, implies that the remaining 10 nm can be attributed to the radial projection of a myosin S2 link.

The protein arrays generating force in isometrically contracting muscle fibers are significantly perturbed by rapid changes in sarcomere length, causing some of the myosin heads to be forcibly detached from actin because of their inability to keep up with the large and fast motions involved. However, the fraction of heads detaching during a quick release has never been described quantitatively. The analysis of stiffness changes after quick release (Ford *et al*., 1985) suggested that a large part of the decrease in T_2_ tension compared to T_0_ is due to reduction in the number of attached myosin heads and partly due as well to a decrease in the average force per cross-bridge. In the present analysis, the latter assumption is unnecessary, and the drop in tension is fully attributed to the decrease in the number of attached myosin heads. The mathematical model describing the mechanics of myosin head detachment was used here to simulate head detachment in both T_1_ and T_2_ phases. The effect of considering myosin head detachment on the relative intensity plot was negligible for small length steps, but gave rise to the characteristic curvature (Huxley & Simmons, 1970) in the length step range 7.0-10.0 nm.

The robustness of the fit to these data in terms of the actin power-stroke model arises from certain features that can be expressed with mathematical rigor with reasonable choices for the free parameters. The consequence of considering these features is a significantly smaller axial movement of myosin heads during all phases of tension recovery compared with predictions of the rotating cross-bridge theory, which are about twice the measured values (Piazzesi et al., 2002). The first feature reducing axial motions of attached myosin heads is that thick and thin filaments can slide relative to each other as a result of structural transitions in actin. When a significant (~30%) fraction of the shortening during a release is attributed to structural changes in actin, axial motions of myosin heads during the elastic response to the length step are correspondingly reduced. The effect on the relative intensity plots for the higher and lower angle peaks of the M3 reflection is a shift upwards of both the T_1_ and T_2_ curves towards the pre-release value as expected.

The second feature that tends to reduce axial motions of cross-bridges is that part of the (linear) strain caused by attached actin filaments undergoing ribbon-to-helix transitions is taken up in azimuthal motions of attached myosin heads (sheer stress). This effect is due to the rotation of ribbon segments about their axes by an angle that is proportional to linear shortening in actin subunits. The net effect is that myosin heads take up strain with rather minimal axial displacements. In particular, torsional resistance gives rise to a significant reduction in S_R2_, the linear shortening due to ribbon-to-helix transitions during the second phase of tension recovery. The overall effect on the relative intensity plots is a slight drop of the T_2_ curve below the T_1_ curve, rather than running systematically above the T_1_ curve. These conclusions are consistent with a careful analysis of the width of equatorial reflections (Huxley *et al*., 1983), which suggested that lateral motions of the myosin heads occur during a contraction, as long as the centers of mass of the heads stay at approximately the same radius, and supported by the estimate sof Reconditi et al. (2004) that the disorder in force-generating myosin heads is +/-17°.

## Conclusion

X-ray interference measurements of axial myosin head movements during transient tension recovery in skeletal muscle fibers by Piazzesi et al. (2002) place severe limits on the range of motions that can be assumed in modeling the mechanics of the contraction process. The results of these experiments present challenges to the classical rotating cross-bridge theory, where the motor element is situated solely in the myosin head itself, because the fit to the experimental curves systematically deviates from the observations (Figure 1). More recent applications of this method (Reconditi et al, 2011) and the discovery that myosin heads in resting muscle are folded against the thick filament backbone (Woodhead et al., 2005; Zoghbi et al., 2008) also raise serious questions about the validity of the widely-accepted role of tropomyosin in regulating the attachment of myosin heads following a tetanus, since the observed movements following activation of troponin by Ca^++^ are simply too fast to be accounted for by current models (Piassezi et al, 2011).

The unexpected behavior of higher orders of the myosin x-ray spacings (~14.5 nm) such as the M6 (Huxley et al., 2006), led to the conclusion that contributions from the thick filament shaft itself are required in order to account for the fine structure of this reflection. However, the contribution to the mass by commensurate actin ribbons has never been considered as the ‘missing’ contribution to the 14.5 nm reflection. The difficulty in explaining the behavior of the M6 (7.15 nm) with the model for myosin head movements derived from fitting the M3 (14.5 nm) curves harkens back to the situation where the 5.9 nm actin-based reflection increases by 31% during a tetanic contraction but *does not shift towards the meridian* as would be expected if myosin heads bound as in the rigor state (Matsubara et al., 1984). We propose that myosin heads ‘mark’ the actin subunits in rigor by adding electron density at higher radius, thus shifting the peak to a lower angle in reciprocal space (Yagi, 1996). In the present case, the ratio of intensities (I_HA_/I_LA_) of higher- and lower-angle peaks of the M6 reflection, problematic for the cross-bridge theory (Huxley et al., 2006), would seem to indicate that actin ribbons, with a 21-screw axis of 7.15 nm, are ‘marking’ the myosin thick filaments, shifting the intensity towards the meridian. The ‘extra density’ on myosin seen in the Fourier maps of Knupp at al. (2009) would seem to support this conclusion.

The actin power-stroke theory, sometimes referred to as the Schutt-Lindberg theory, gives an excellent fit to the x-ray interference measurements (as shown in Figures 7–9) of Piazzesi et al. (2002). The classical equation (Huxley & Simmons,1971) describing tension recovery following a rapid drop in tension, a foundation of the myosin cross-bridge theory, has been shown here to be a simple consequence of cooperative structural transitions along actin filaments, regulating the attachment and detachment of myosin heads. A re-interpretation of force-energy data obtained from *in vitro* motility assays using isolated actin and myosin (Schutt & Lindberg, 1998) leads to an equally simple derivation of the classical Hill equation (Hill, 1938), and a resolution of many of the problems posed by *in vitro* studies. These results should compel a serious consideration of the concept of actin as the independent force generator in contracting muscle fibers (Schutt & Lindberg, 1992;1998).

## Supporting information

Supplemental videos for Schutt et al. - Mucle Contraction as a Markov Process - II

## APPENDIX A

### Simulation of the X-ray diffraction pattern

A program was written in C to simulate the X-ray diffraction pattern based on our model. In the main execution, point scatterers were located at the centers of mass of myosin and actin molecules, with their scattering weights proportional to the molecular weights of the scatterers. Such an approach was the most economical one from the viewpoint of computational time. More precise calculations using the atomic model of myosin (Rayment *et al*., 1993) and actin ribbons (Schutt & Lindberg, 1993) were performed occasionally to check whether the point-scatterer approach was valid, which was found to be true. We placed four ribbon segments (each 42.9 nm long) in the overlap zone of every actin filament, randomly selecting their position. As a result, at every myosin head layer in the overlap zone, there appeared a corresponding population of attached myosin heads across the fiber.

The program determined the number of myosin head layers in the overlap zone, taking into account the lengths of actin filaments, myosin filaments, sarcomere length and the spacing between the myosin head layers. The program called a subroutine (GenerateEvenSpacings) to create an array of coordinates with uniform spacing of 14.57 nm representing the centers of mass of the myosin heads. The “mirror” array was also generated, which corresponds to the myosin heads on the opposite side of the bare zone. The outer program loop ran through the values of the release length step from 0 Å to 10.0 nm with an increment of 0.1 nm. For each value of the length step, a subroutine (QuickRelease) was called, which performed calculations necessary for computing the perturbed axial coordinates of the myosin heads during the T_1_ and T_2_ phases. These coordinates were then supplied to the internal loop, which covers the range of reciprocal lengths from 1/16.6 nm^-1^ to 1/12.5 nm^-1^ with an increment of 10^-6^ nm^-1^, giving rise to 20,000 simulated data points in reciprocal space. In order to simulate each data point, a subroutine (FourierTransform) was called, which calculates the intensity of the corresponding X-ray reflection via the Fourier transform. Finally, a subroutine (AnalyzePeaks) was called, which returns the the simulated diffraction pattern in the vicinity of the expected M3 reflection (1/14.5 nm^-1^) and extracts information about the intensities and spacings of the higher- and lower-angle peaks of the simulated M3 reflection.

### Computation of myosin head axial positions

The most important block of the simulation program is a subroutine (QuickRelease) that calculates T_1_ and T_2_ tensions, probabilities of myosin head detachment, and the axial positions of attached and detached heads.

The 5.1 nm/T_0_ length-tension relationship (Piazzesi *et al*., 2002) was used to extract the extreme tension drop during the T_1_ phase. We calculated myosin head compliance through equations (4,5,6) and the probabilities of myosin head detachment in phases T_1_ and T_2_ through equations (14,15,16,17). The functions (equation 19 and 23) allowed the T_2_ tension to be estimated with equation (18).

In order to calculate the shift of axial position of a myosin head, two elementary shifts were calculated: Δ*x_a_*, which is the shift of the actin monomer that is bound to the myosin head, and Δ*x_m_*, which is the shift of the point where the myosin S2 rod is connected to the myosin filament. These shifts varied from head to head because of the distribution of tension in the filaments, and each of the shifts was a function of the initial axial coordinate *z* of the head. In the T_1_ phase, the tension decreases from T_0_ to T_1_, so that:

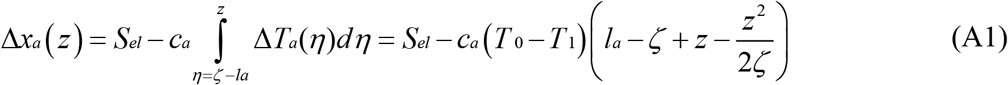

and similarly:

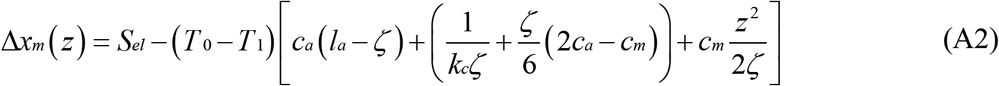

The change of strain of a myosin molecule can be found as Δ*x_m_* (*z*) – Δ*x_a_* (*z*). Recalling that the pivot in the myosin head is located in the converter domain (Dobbie *et al*., 1998), two modes of myosin head elasticity (Fig. A-2) and certain weights attributed to them can be calculated. This allows the change of length of the S2 rod, and the shift of the head-rod junction point to be determined. Taking into account that only the myosin light-chain domain is flexible, while the catalytic domain does not move relative to the actin-binding site, and considering that roughly 80% of myosin head’s density is in the catalytic domain, the displacement of center of mass for each myosin head can be calculated during the T_1_ phase.

Calculation of axial positions of myosin heads during the T_2_ phase was more sophisticated. To take account of torsional coupling, the values of S_R2_ were calculated independently for each head (the graph shown in Fig. 5, in fact, had values of S_R2_ averaged over the overlap zone). The algorithm for calculating the myosin head axial positions during the T_2_ phase consisted of two parts. In the first part, structural transitions in actin during the T_2_ phase were temporarily ignored, and elementary myosin and actin shifts Δ*x_a_* 2 and Δ*x_m_* 2 were calculated, relative to their positions during the T_1_ phase, solely as an elastic response to the regenerating tension. The shift of myosin is:

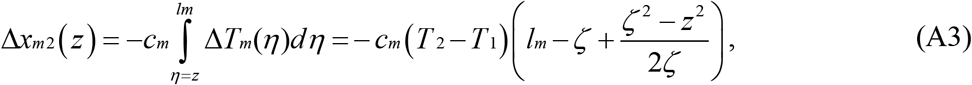

in the direction opposite to its shift during the T_1_ phase; while the shift of actin is:

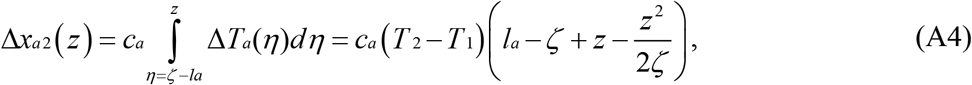

in the same direction (towards the middle of sarcomere) as its shift during the T_1_ phase. The opposite movement of actin is impossible during the T_2_ phase, because the position of Z-line is fixed as soon as the sarcomere length is fixed at a new value. Both Δ*x*_*a*2_ and Δ*x*_*m*2_ tend to decrease the length of S2 links, so the total decrease of S2 length due to elastic response in filaments to regenerating tension is the sum of absolute values of Δ*x*_*a*2_ and Δ*x*_*m*2_. If we consider the actin-binding site fixed, then even further decreases in the length of S2 result from tilting of the myosin light-chain domain, which is the elastic response of myosin head itself to the recovering tension. All these factors allow us to calculate intermediate values for the “compressed” length of S2 rod.

In the second part of the T_2_ calculation, structural transitions in actin are activated, yielding the length S_R2_, by which r-actin contracts during the T_2_ phase. Since the Z-line is fixed, the length S_R2_ will be equal to the shift of myosin-bound r-actin monomer towards the Z-line, and it will effect an increase in S2 length through equations (equations 10–13) that include torsional coupling. Having calculated the “compressed” S2 length, and assuming that actual S2 length after tension recovery is the same as its length before the release, S_R2_ is obtained by solving a non-linear equation.

At this point in the calculation, the coordinates for each myosin head are available corresponding to states: 1) during the extreme drop of tension (extreme T_1_); and 2) at the end of tension recovery (asymptotic T_2_). It is necessary to take into account that, because of finite rate of tension recovery, the measured tension at any moment of time is between T_1_ and T_2_:

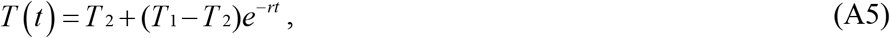

where the rate of recovery is *r* = 0.2* (1 + *e*^0.5*S*^); S is the value of length step, positive for release. The times t_1_ and t_2_ in the actual experiment were 110μs and 2.0 ms, respectively (Piazzesi *et al*., 2002). The time course of myosin heads was taken as a shift from the extreme T_1_ to the asymptotic T_2_ positions during the time course of tension recovery, and thus the axial coordinates of myosin heads through the recovery formula could be calculated through equation (28) for tension recovery:

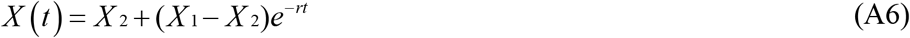

Since a certain fraction of myosin heads that were attached before the quick release detach during the process, the motions of detached heads need to be calculated in order to get an accurate calculation of the X-ray diffraction pattern. Since the detached heads are not bound to actin, they do not bear tension. Therefore, irrespective of the timing at which those heads detach, the springs in the S2 elastic links of these detached heads recoil to a length corresponding to zero tension. This length can be determined, since it is the same as the length of the S2 links in the attached heads at the T_1_ phase of the release step S = 5.1 nm, which was found to bring the tension in the filaments to zero (Piazzesi *et al*., 2002). The angle of tilt of the light-chain domain is, however, likely to vary from one detached head to another, and it would depend on the state of the head immediately before detachment. Therefore, the heads that detach during the quick release are partially disordered. When considering the scattering from a population of such heads, the disorder is taken account of by multiplying the scattering factor by a “disorder factor”, a number between 0 and 1 that is the smaller, the larger the statistical variance in head positions.

### Calculation of actin scattering centers

In calculating the positions of actin scatterers we considered that, during the release, every ribbon segment shifted relative to the myosin head attached to it by the distance L_R_ (equation 15)); however, if a head detaches, the ribbon associated with it is simply pulled by elastic forces following detachment. If we define the position of a ribbon by the axial coordinate *x_R_* of its outside boundary relative to the actin-binding site of the associated myosin head, then, starting from the pre-release ribbon position *x_R_* = –*x* (see Fig. 5), we can calculate its shifted position 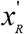 as:

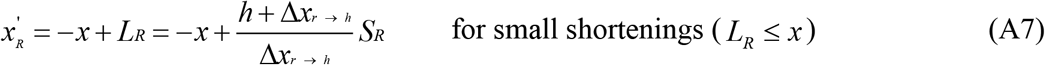

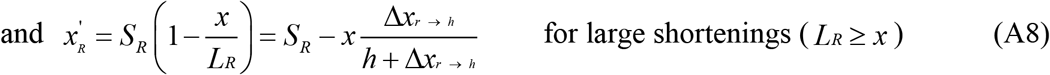

In the case of *L_R_ ≥ x* (detachment case), the outside ribbon boundary overshoots past a myosin head by a distance equal to the distance of filament sliding (S_R_) minus a portion of S_R_ that was necessary to knock the head off. Both formulae (A7) and (A8) give 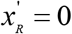 at the shortening corresponding to the myosin head detachment (*L_R_ = x*).

In its simplest form, the actin power-stroke theory has only one myosin head attached for every ribbon segment. During an isometric contraction there are, on average, four ribbon segments per thin filament at full overlap since the force generated by each contracting ribbon segment is approximately one quarter of the maximum isometric tension (Schutt & Lindberg, 1998). It follows (considering 1:2 myosin:actin ratio in the lattice) that about 8 myosin layers in a given thick filament are bound to a given actin filament at any moment. Since there are around 50 layers of myosin heads in the overlap zone at full overlap, it means that only about 15% of myosin heads are attached. This fraction of attached myosin heads is considerably smaller than the corresponding values of around 50% (Linari *et al*., 1998), assumed in most treatments based on the cross-bridge theory.

Thus, the scattering pattern results from a relatively small number of myosin heads. This pattern will depend on whether, at any given time, attached myosin heads appear in all (~50) layers of myosin heads or in 8 layers only. The first possibility will be true if the ribbon segments are located in random places of different thin filaments within a sarcomere. If, on the other hand, the ribbon segments in the individual thin filaments are strictly in register, we will have a set of discrete disks across the sarcomere in which the diffracting myosin heads will appear. The original version of the actin power-stroke theory did not elaborate on whether the ribbons are in register or whether transitions in different actin filaments are coupled to each other at all. However, simulation of the M3 X-ray reflection pattern with the “discrete disk” model showed a triplet of doublets, with two sets of side peaks considerably far away from the main reflection, with the spacing between the main doublet and the satellites being the reciprocal of the sampling distance between the disks (Fig. 8). The side peaks of such intensities and in such positions have never been observed experimentally. The model in which the ribbons were considered to populate the overlap zone in a random fashion has, on the other hand, gave rise to one doublet, which was at the correct position. This rather simple modeling experiment argues very strongly in support of the idea that the positions of ribbon segments in various actin filaments are not correlated to each other, so that ribbons, at a given instant of time during an isometric contraction, can be considered randomly distributed along the overlap zone (see Schutt and Lindberg, 1998).

## SUPPLEMENTARY MATERIAL

### Mechanical description of muscle contraction

#### Tension distribution

The thick and thin muscle filaments and myosin cross-bridges in a sarcomere are interconnected in a mechanically complex series-parallel arrangement. The elements of a sarcomere that are mechanically in series are the Z-line, the zone of non-overlapped actin filaments, the zone of actin and myosin filament overlap, and the bare zone of myosin filaments. In the overlap zone, the cross-bridges link actin and myosin filaments and bear tension. During a contraction, the tension T_0_ is fully borne by actin filaments in the I-band, and is distributed between actin and myosin filaments via cross-bridges in the overlap region. Tension is fully borne by the myosin filament in the bare zone. The assumption that cross-bridges populate the overlap zone uniformly, and are equivalent in the sense of all having the same stiffness, enables the distribution of tension along the actin and myosin filaments in the overlap region to be expressed in a rather simple way. If ζ is the length of the overlap area, and *z* is the axial coordinate of the point in question, counting from the outside boundary of the overlap area, then the tensions Ta in actin filament and Tm in myosin filament depend linearly on this coordinate in the overlap area (Fig. SM-1):

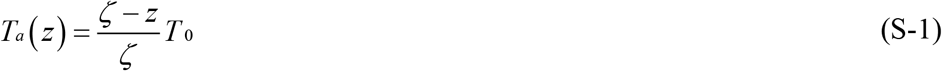

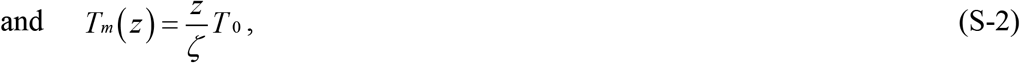

where the overlap area is defined by *z* being within the range 0 ≤ *z ≤ζ*.

**FIGURE SM-1.**
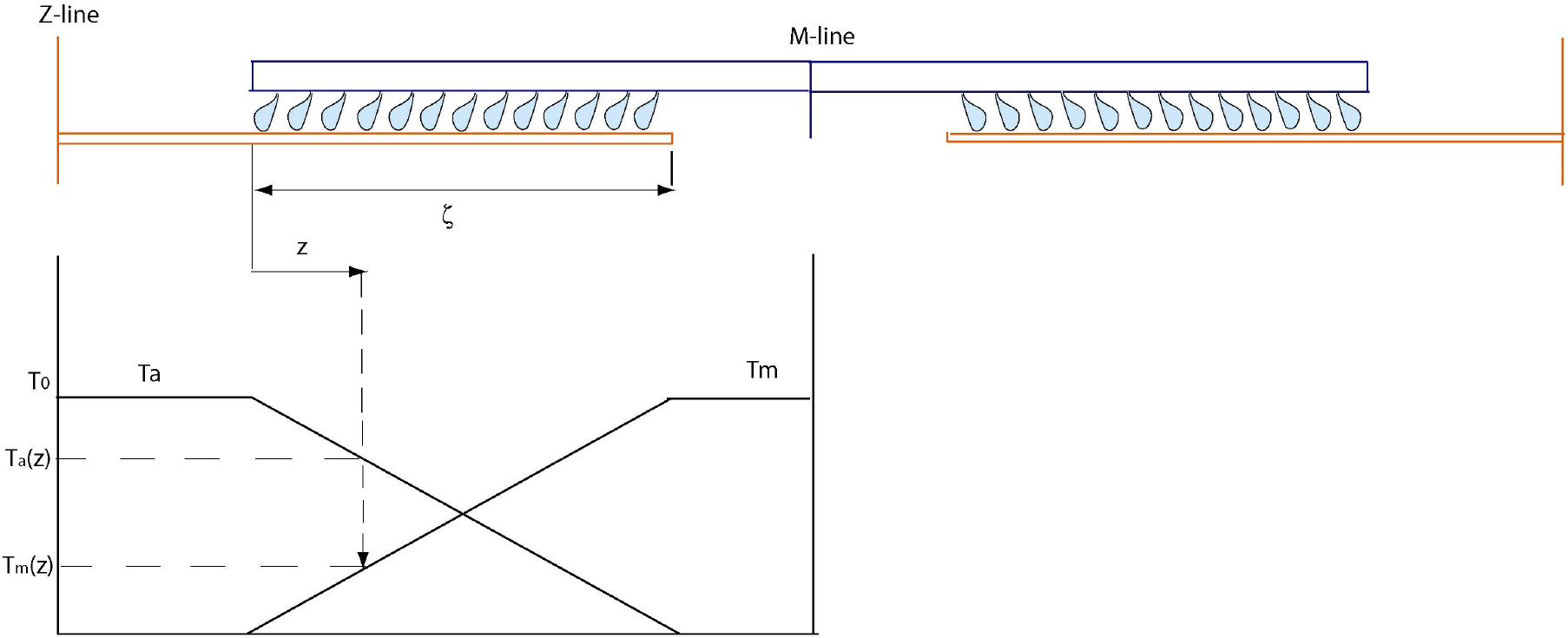
Variables used to distribute tension between actin and myosin filaments.

#### Filament compliance

A convenient parameter for describing the elasticity of a mechanical element is compliance. If a change in applied force ΔF causes the element of length x to elongate by Δx, then the stiffness of the element is:

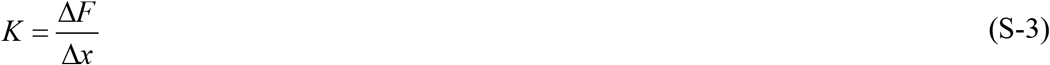

and the compliance (an inverse of stiffness) is:

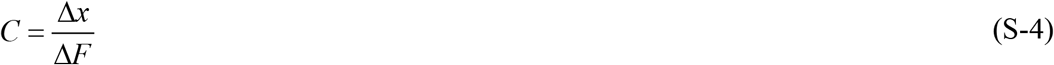

In the field of muscle contraction, it is convenient to scale compliances to tension (which is force per cross-sectional area). Also, a frequently used variable is compliance of a filament per unit length. It shows the fractional elongation of a filament due to a given change ΔT in tension:

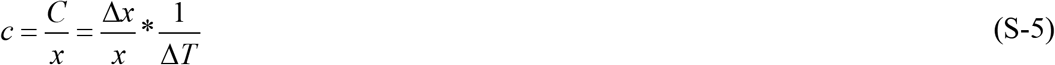

One of the methods of obtaining compliances of actin and myosin filaments is to estimate them from the values of average strain in the filaments. The strains in actin and myosin filaments are experimentally deduced from the relative changes in spacings of X-ray reflections during a muscle stretch (H.E. Huxley *et al*., 1994; Wakabayashi *et al*., 1994) and were reported as 0.12±0.01%/T_0_ and 0.23±0.01%/T_0_ for myosin and actin filaments respectively. For the myosin filament, the experimentally observed strain is averaged over the overlap zone only, since the myosin heads that contribute to the myosin X-ray reflections are absent in the bare zone. The relationships between the filament compliances and the average strains are derived (Linari *et al*., 1998) with the consideration of the tension distribution along the filaments:

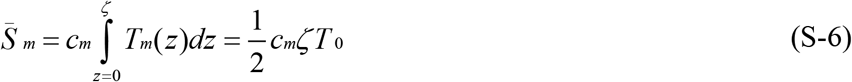

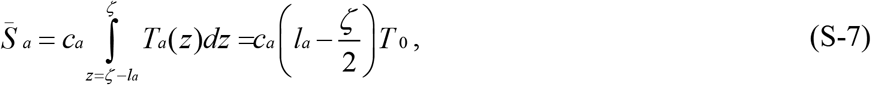

where *l_a_* is length of an actin filament.

An alternative experimental method of studying the compliance of the actin filaments takes advantage of the fact that in the sarcomere length range 2.00 – 2.17 μm the length of the overlap zone can be considered constant. Therefore, as the sarcomere length is increased, the I-band length increases by the same amount. The variation in the total half-sarcomere compliance would be then solely due to the variation in the I-band compliance, which is equal to actin compliance times the I-band length. It follows that if s_0_ is the reference sarcomere length, and C_0_ is the half-sarcomere compliance at the reference sarcomere length, then the half-sarcomere compliance varies linearly with sarcomere length:

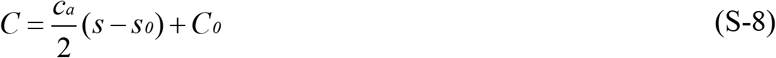

Actin compliance is then deduced as the slope of this relationship. This experiment (Linari *et al*., 1998) showed the value of actin compliance equal to 2.32±0.34 nm μm^-1^/T_0_, corresponding to the average strain in actin filament of 0.21±0.03%/T_0_.

#### Myosin head compliance

The existence of instantaneous compliance within each myosin head is a key element for the ability of sarcomeres to shorten during a quick release (Huxley & Simmons, 1971). As the strain in the myosin head is relieved, actin and myosin filaments can slide relative to each other. Originally, the myosin head elastic element was attributed to the S2 subfragment that links a myosin head to the body of myosin filament (Huxley, 1969; Huxley & Simmons, 1971). The S2 link is predominantly a-helical and can be treated as a spring, the extension of which is proportional to tension through Hooke’s law.

It was shown later (Dobbie *et al*., 1998) that the myosin head light chain domain has the ability to bend elastically relative to the actin-bound catalytic domain, which also gives rise to the filament sliding. The bending of the light chain domain is accompanied by a local conformational change in the converter domain, which is located at the junction of the catalytic and light chain domains; but the rest of the density of the catalytic domain does not move relative to the actin-binding site. The length by which filaments slide as a result of the change of the tilt of the light-chain domain by angle a is *L* sin*α*, where L is the length of the light-chain domain (~9.5 nm). We will further consider the myosin head compliance distributed between the two elastic modes, one of which corresponds to stretching of the S2 link, and the other one to tilting of the light chain domain. Both these modes are schematically demonstrated in Fig. SM-2.

**FIGURE SM-2.**
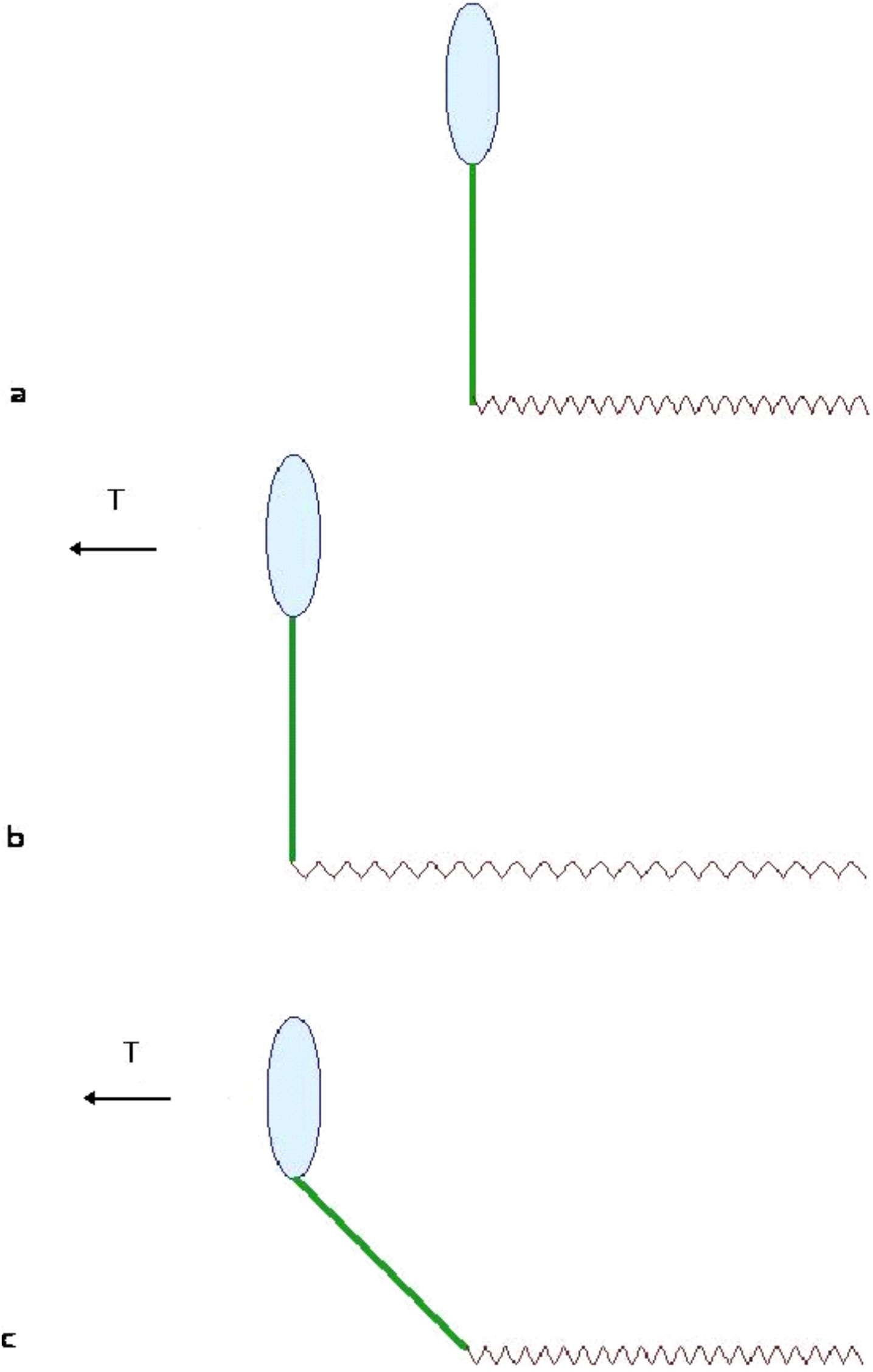
Illustration of bending and stretching modes in myosin used in the calculation of myosin head compliances.

#### Half sarcomere compliance

Because of the parallel/series arrangement of mechanical elements in a sarcomere, the problem arises of how to calculate half-sarcomere compliance, that is, the extension of one half-sarcomere as a response to a unit change in tension. This problem has been studied (Ford *et al*., 1981), and the formula derived is:

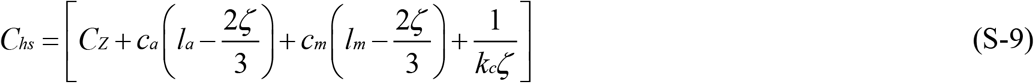

Here ζ is the length of overlap zone, l_a_ – length of actin filament, l_m_ – length of myosin filament (corresponding to one of the sides of the M-line), c_a_ and c_m_ – actin and myosin filament compliances (per unit length), k_c_-myosin head stiffness per unit length, C_Z_-compliance of the Z-line.

The basic meaning of the equation (3.9) is that the total extension of the half-sarcomere is the sum of elastic extensions in the Z-line, the actin filament, the myosin filament, and the sliding distance between the actin and myosin filaments due to myosin head compliance. Equation (S-9) for total half-sarcomere compliance is widely used in the modern literature on muscle mechanics (Linari *et al*., 1998, Piazzesi *et al*., 2002), as it provides a method of estimating myosin head compliance, which is not measurable experimentally. The total half-sarcomere compliance is known from the length-tension relationship, and the contributions to the half-sarcomere shortening from filament elasticities can be calculated from filament compliances. In the absence of evidence for compliance of the Z-line, the term C_Z_ is normally assumed to be zero (Linari *et al*., 1998); so that the remaining term, which is the myosin head compliance, can be readily extracted.

We have already seen how the extensions in actin and myosin filaments are related to tension. So, where do coefficients of the “2ζ/3” type come from in the equation (S-9)? Qualitatively, they come from allowing filaments to slide not only due to myosin head compliance, but also as a result of the difference between the actin and myosin filament compliances. We have made a more detailed analysis of the underlying mathematical section (Ford *et al*., 1981) to find out how specific fragments of the half-sarcomere would extend due to a unit change in tension. Referring to Fig. SM-3 for a definition of points A, B, C, D, E and F, it follows that the lengths of the filament segments AB, BD, CE, EF and the axial projections of cross-bridge strains BC and DE on the opposite sides of the overlap zone change the following way owing to the change Δ*T* in tension:

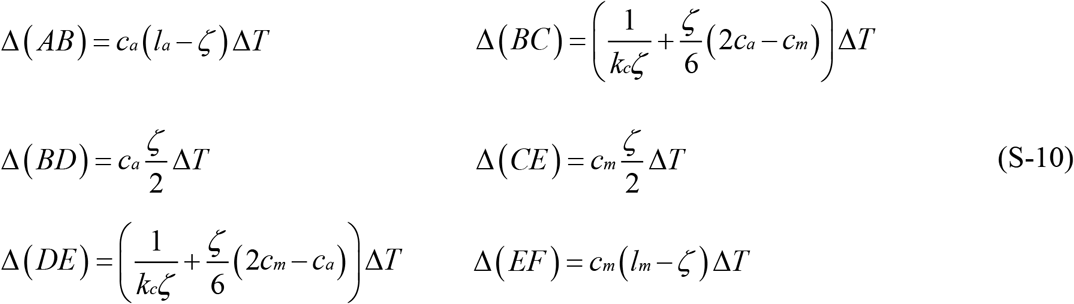

Summation of individual strains from the point A to the point F (by following either of the paths) gives the total extension of a half-sarcomere, which is equal to the half-sarcomere compliance times tension:

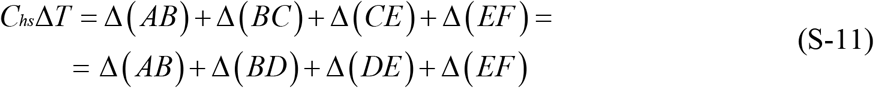

**FIGURE SM-3.**
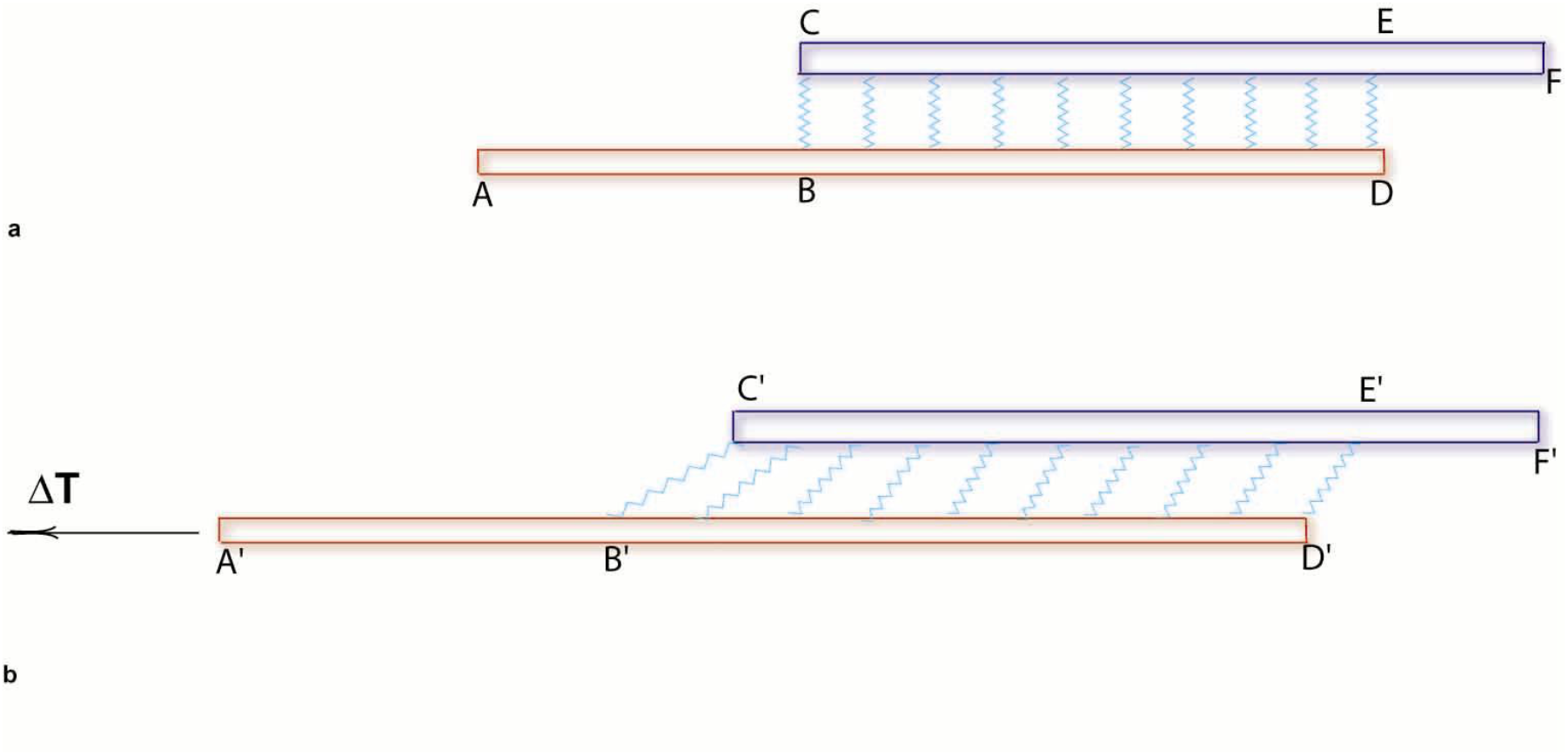
Definition of coordinates used to calculated the effect of different values for the compliance in actin and myosin thick filaments.

## Notes

### Competing Interest Statement

The authors have declared no competing interest.

## References

Brenner, B. (1991) Rapid dissociation and re-association of acto-myosin cross-bridges during force generation: a newly observed facet of cross-bridge action in muscle. Proc. Natl. Acad. Sci. USA 88, 10490–10494.

Brunello, E., Bianco, B., Piazzesi, G., LInari, M., Reconditi, M., Panine, P., Narayanan, T., Helsby, W.I., Irving,, M., and Lomardi, V. (2006) Structural changes in the myosin filament and cross-bridges during active force development in single intact frog muscle fibres: stiffness and X-ray diffraction measurements. J. Physiol. 577, 971–984.

Brunello, E., Caremani, M., Melli, L., Linari, M., Fernandez-Martinez, M., Narayanan, T., Irving, M., Piazzesi, G., Lombardi, V. and Reconditi, M. (2014) The contribution of filament and cross-bridges to sarcomere compliance in skeletal muscle. J. Physiol. 592, 3881–3899.

Cecchi, G., Griffiths, P.J., and Taylor, S. (1982) Muscular contraction: kinetics of cross-bridge attachment studied by high-frequency stiffness measurements. Science 217, 70–72.

Chen, Y.-D., and Brenner, B. (1993) On the regeneration of the actin-myosin power stroke in contracting muscle. Proc. Natl. Acad. Sci USA 90, 5148–5152.

Chik, J.K., Lindberg, U., and Schutt, C.E. (1996) The structure of an open state of ß-actin at 2.65 Å resolution. J. Mol. Biol. 263, 607–623.

Cooke, R. (1986) The mechanism of muscle contraction. CRC Crit. Rev. Biochem. 21, 53–118.

Dobbie, I., Linari, M., Piazzesi, G., Reconditi, M., Koubassova, N., Ferenczi, M.A., Lombardi, V., and Irving, M. (1998) Elastic bending and active tilting of myosin heads during muscle contraction. Nature 396, 383–387.

Eisenberg, E., Hill, T.L., and Chen, Y.-D. (1980) Cross-bridge model of muscle contraction. Biophys. J. 29, 195–227.

Ford, L.E., Huxley, A.F., and Simmons, R.M. (1977) Tension responses to sudden length change in stimulated frog muscle fibres near slack length. J. Physiol. (Lond.) 269, 441–515.

Ford, L.E., Huxley, A.F., and Simmons, R.M. (1981) The relation between stiffness and filament overlap in stimulated frog muscle fibres. J. Physiol. (Lond.) 3ll, 219–249.

Ford, L.E., Huxley, A.F., and Simmons, R.M. (1985) Tension transients during steady shortening of frog muscle fibres. J. Physiol. (Lond.) 361, 131–150.

Geeves, M.A., and Holmes, K.C. (1999) Structural mechanism of muscle contraction. Annu. Rev. Biochem. 68, 687–728.

Harada, Y., Sakurada, K., Aoki, T., Thomas, D.D., and Yanagida, T. (1990) Mechanochemical coupling in actomyosin energy transduction studied by *in vitro* movement assay. J. Mol. Biol. 216, 49–68.

Higuchi, H., and Goldman, Y.E. (1991) Sliding distance between actin and myosin filaments per ATP molecule hydrolysed in skinned muscle fibres. Nature 352, 352–354.

Higuchi, H., Yanagida, T., and Goldman, Y.E. (1995) Compliance of thin filaments in skinned fibers of rabbit skeletal muscle. Biophys. J. 69, 1000–1010.

Higuchi, H., and Goldman, Y.E. (1995) Sliding distance per ATP molecule hydrolyzed by myosin heads during isotonic shortening of skinned muscle fibers. Biophys. J. 69, 1491–1507.

Hill, A.V. (1938) The heat of shortening and the dynamic constants of muscle. Proc. Roy. Soc. Lond. B126, 136–195.

Holmes, K.C. (1997) The swinging lever-arm hypothesis of muscle contraction. Curr. Biol. 7, 112–118.

Huxley, A.F. (1980) Reflections on Muscle Contraction, The Sherrington Lectures, Princeton Univ. Press, Princeton, NJ.

Huxley, A.F., and Simmons, R.M. (1971) Proposed mechanism of force generation in striated muscle. Nature 233, 533–538.

Huxley, H.E., and Brown, W. (1967) The low-angle x-ray diagram of vertebrate striated muscle and its behaviour during contraction and rigor. J. Mol. Biol. 30, 383–434.

Huxley, H.E. (1969) The mechanism of muscular contraction. Science, 164, 1356–1366.

Huxley, H., Reconditi, M., Stewart, A., and Irving, T. (2006) X-ray interference studies of cross-bridge action in muscle contraction: evidence from quick releases. J.Mol.Biol., 363, 743–761

Huxley, H.E., Simmons, R.M., Faruqi, A.R., Kress, M., Bordas, J., and Koch, M.H.J. (1983) Changes in the X-ray reflections from contracting muscle during rapid mechanical transients and their structural implications. J. Mol. Biol. 169, 469–506.

Huxley, H.E., Stewart, A., Sosa, H., and Irving, T. (1994) X-ray diffraction measurements of the extensibility of actin and myosin filaments in contracting muscle. Biophys. J. 67, 2411–2421.

Irving, M., Lombardi, V., Piazzesi, G., and Ferenczi, M.A. (1992) Myosin head movements are synchronous with the elementary force-generating process in muscle. Nature 357, 156–158.

Juanhuix, J., Bordas, J., Campmany, J., Svensson, A., Bassford, M.L., and Narayanan, T. (2001) Axial disposition of myosin heads in isometrically contracting muscles. Biophys. J. 80, 1429–1441.

Knupp, C., Offer, G., Ratatunga, K.W., and Squire, J.M. (2009) Probing Muscle Myosin Motor Action: X-Ray (M3 and M6) Interference Measurements Report Motor Domain Not Lever Arm Movement. J. Mol. Biol. 390, 168–181.

Knupp, C. and Squire, J.M. (2019) Myosin Cross-bridge Behaviour in Contracting Muscle – The T1 Curve of Huxley and Simmons (1971) Revisited. Int J. Mol. Sci. 20, 4892–4924.

Linari, M., Dobbie, I., Reconditi, M., Koubassova, N., Irving, M., Piazzesi, G., and Lombardi, V. (1998) The stiffness of skeletal muscle in isometric contraction and rigor: the fraction of myosin heads bound to actin. Biophys. J. 74, 2459–2473.

Linari, M., Piazzesi, G., Dobbie, I., Koubassova, N., Reconditi, M., Narayanan, T., Diat, O., Irving, M., and Lombardi, V. (2000) Interference fine structure and sarcomere length dependence of the axial x-ray pattern from active single muscle fibers. Proc. Natl. Acad. Sci. USA 97, 7226–7231.

Lombardi, V., Piazzesi, G., and Linari, M. (1992) Rapid regeneration of the actin-myosin power stroke in contracting muscle. Nature 355, 638–641.

Lombardi, V., Piazzesi, G., Ferenczi, M.A., Thirlwell, H., Dobbie, I., and Irving, M. (1995) Elastic distortion of myosin heads and repriming of the working stroke in muscle. Nature 374, 553–555.

Lymn, R.W., and Taylor, E.W. (1971) Mechanism of adenosine triphosphate hydrolysis by actomyosin. Biochemistry 10, 4617–4624.

Månsson, A., Rassier, D., and Tsiavaliaris, G. “Poorly Understood Aspects of Striated Muscle Contraction,” BioMed Research International, vol. 2015, Article ID 245154, 28 pages, 2015. doi:10.1155/2015/245154

Matsubara, I., Yagi, N., Miura, H., Ozeki, M. and Isumi, T. (1984) Intensification of the 5.9 nm actin layer line in contracting muscle. Nature 312, 471–473.

Morel, J.E. (2016) Molecular and Physiological Mechanisms of Muscle Contraction. CRC Press, Boca Raton, FL.

Piazzesi, G., Reconditi, M., Linari, M., Lucii, L., Sun, Y.B., Narayanan, T., Boesecke, P., Lombardi, V., and Irving, M. (2002) Mechanism of force generation by myosin heads in skeletal muscle. Nature 415, 659–662.

Podolsky, R.J. (1960) Kinetics of muscular contraction: the approach to steady state. Nature 188, 666–668.

Podolsky, R.J., Nolan, A.C., and Zaveler, S.A. (1969) Cross-bridge properties derived from muscle isotonic velocity transients. Proc. US. Nat. Acad. Sci. 64, 504–511.

Pollack, G.H. (1990), Muscles & Molecules: Uncovering the Principles of Biological Motion, Ebner & Sons, Seattle, WA

Reconditi, M., Linari, M., Lucii, L., Stewart, A., Sun, Y-B, Boesecki, P., Narayanan, T., Fischetti, R.F., Irving, T., Piazzesis, G., Irving, M. and Lombardi, V. (2004) The myosin motor in muscle generates a smaller and slower working stroke at higher load. Nature 428, 578–581

Reconditi, M., Brunello, E., Linari, M., Bianco, P., Narayanan, T., Panine, P., Piazzesi, G., Lombardi, V., and Irving, M. (2011) Motion of myosin head domains during activation and force development in skeletal muscle. Proc. Natl. Acad. Sci. USA 108, 7236–7240.

Schutt, C.E., and Lindberg, U. (1992) Actin as the generator of tension during muscle contraction. Proc. Natl. Acad. Sci. USA 89, 319–323.

Schutt, C.E., and Lindberg, U. (1993) A new perspective on muscle contraction. FEBS Lett. 325, 59–62.

Schutt, C.E., and Lindberg, U. (1998) Muscle contraction as a Markov process I: energetics of the process. Acta Physiol. Scand. 163, 307–323.

Schutt, C.E., Lindberg, U., Myslik, J. and Strauss, N. (1989) Molecular packing in profilin:actin crystals and its implications, J. Mol. Biol. 209, 735–746.

Schutt, C.E., Myslik, J.C., Rozycki, M.D., Goonesekere, N.C., and Lindberg, U. (1993) The structure of crystalline profilin-ß-actin. Nature 365, 810–816.

Schutt, C.E., Rozycki, M., Chik, J.K, and Lindberg, U. (1995a) Structural studies on the ribbon-to-helix transition in Profilin:Actin crystals. Biophys. J. 68, 12s–18s.

Toyoshima, Y.Y., Kron, S.J., and Spudich, J.A. (1990) The myosin step size: measurement of the unit displacement per ATP hydrolyzed in an *in vitro* assay. Proc. Natl. Acad. Sci. USA 87, 7130–7134.

Uyeda, T.Q., Kron, S.J., and Spudich, J.A. (1990) Myosin step size. Estimation from slow sliding movement of actin over low densities of heavy meromyosin. J. Mol. Biol. 214, 699–710.

Wakabayashi, K., Sugimoto, Y., Tanaka, H., Ueno, Y., Takezawa, Y., and Anemiya, Y. (1994) X-ray diffraction evidence for the extensibility of actin and myosin filaments during muscle contraction. Biophys. J. 67, 2422–2435.

Woodhead, J.L., et al. (2005) Atomic model of a myosin filament in the relaxed state. Nature 436:1195–1199.

Zoghbi, M.E., Woodhead, J.L., Moss, R.L., and Craig, R. (2008) Three-dimensional structure of vertebrate cardiac muscle myosin filaments. Proc. Natl. Acad. Sci. USA 105, 2386–2390.

Xu, S.G., Kress, M., and Huxley, H.E. (1987) X-ray diffraction studies of the structural state of crossbridges in skinned frog sartorius muscle at low ionic strength. J. Muscle Res. Cell Motil. 8, 39–54.

