## Supplementary figures and images for "Muscle contraction as a Markov Process - II: x-ray interference data (M3 & M6 reflections) imply myosin cross-bridge motions are controlled by structural transitions along actin filaments"

### Supplemental videos for Schutt et al. - Mucle Contraction as a Markov Process - II

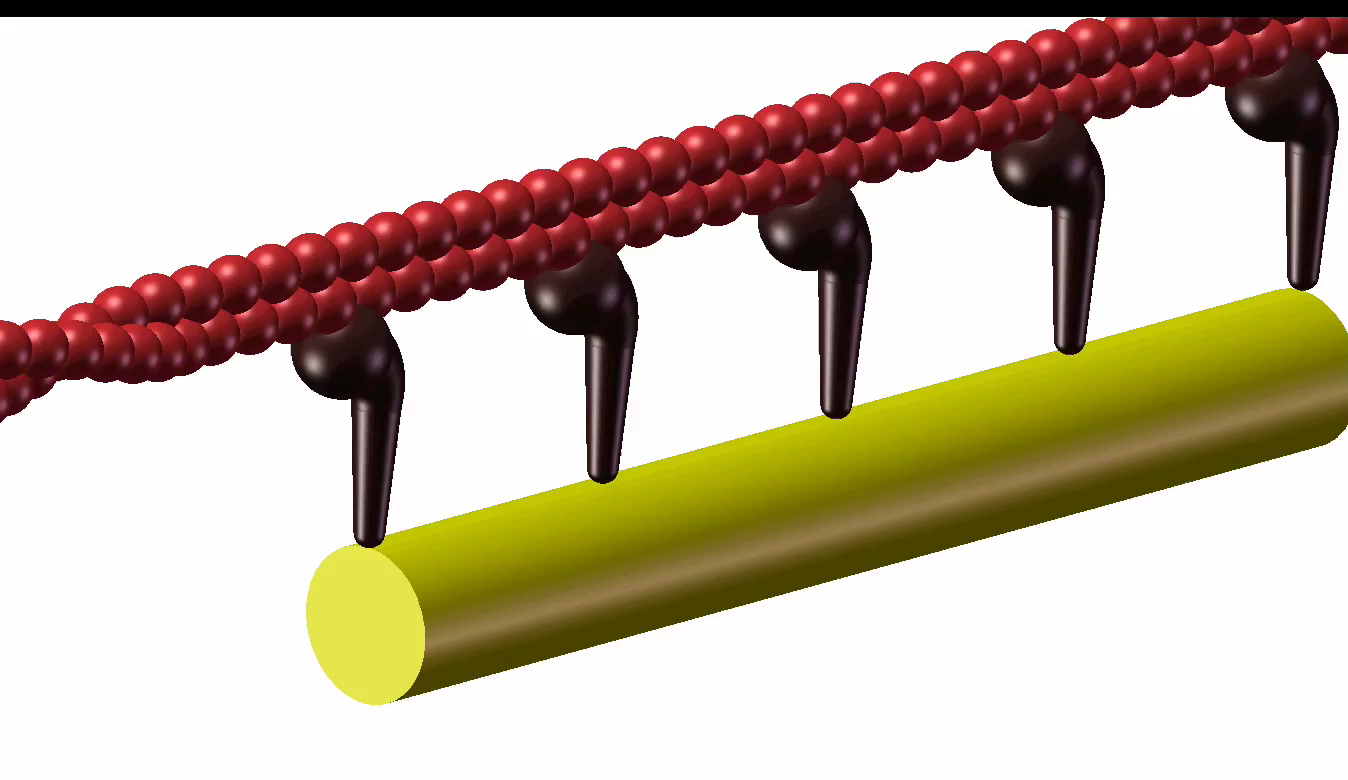

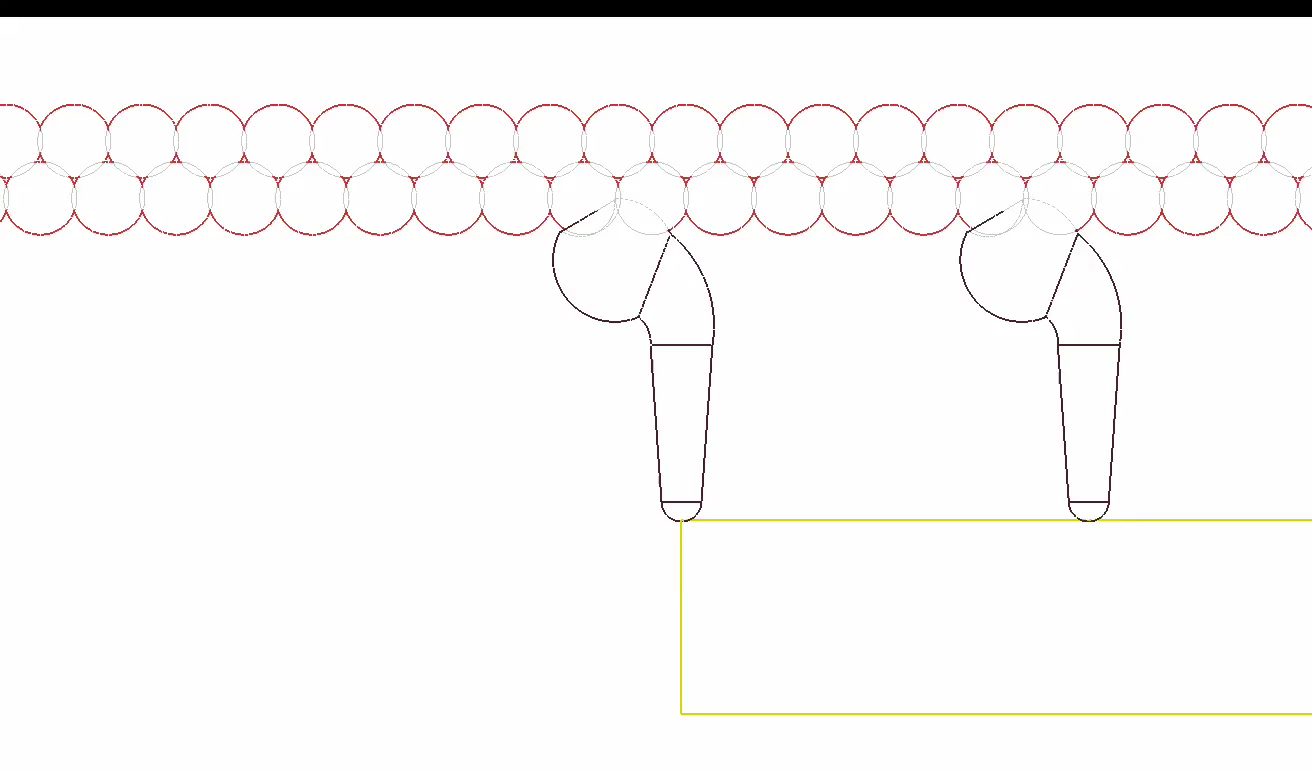

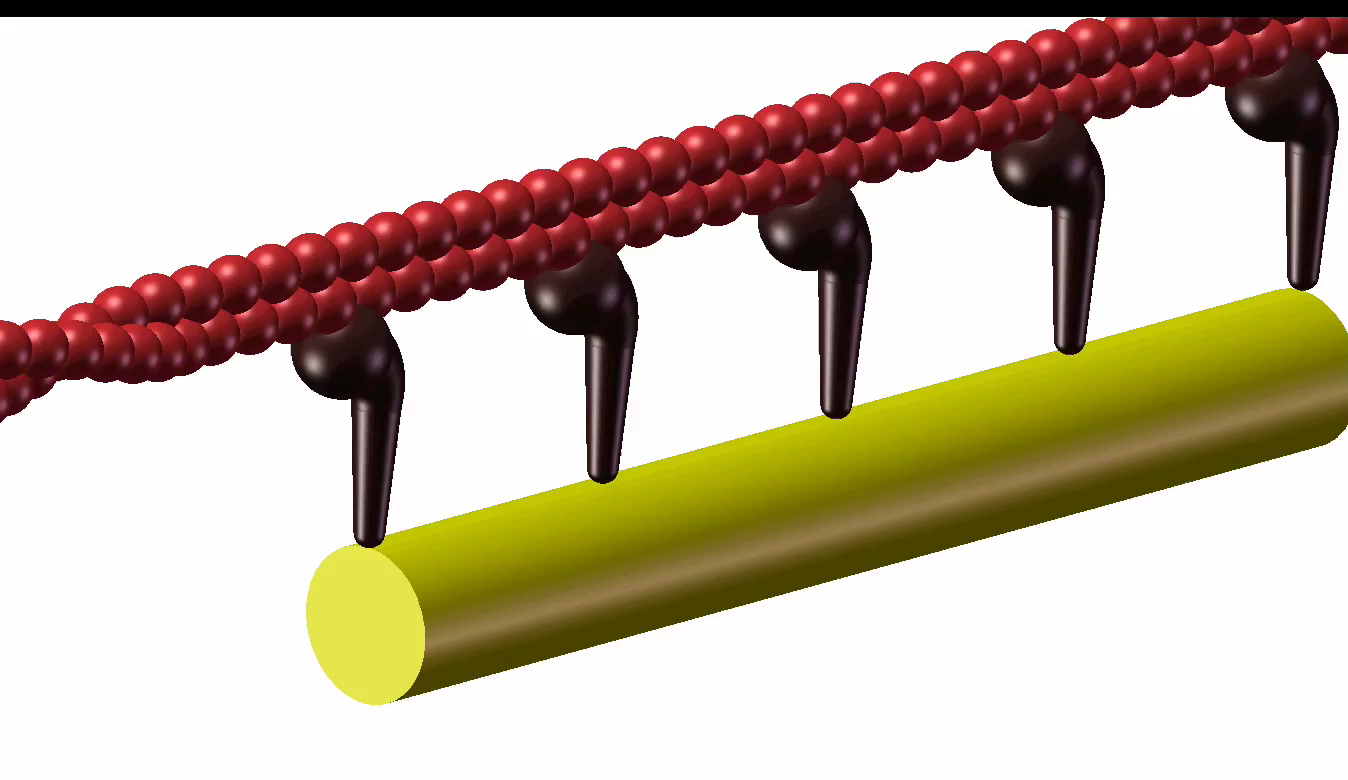
